# Δ^9^-tetrahydrocannabinol (THC) Increases the Rewarding Value of Oxycodone During Self-Administration in Rats

**DOI:** 10.1101/2024.01.18.576277

**Authors:** Jacques D. Nguyen, Yanabel Grant, Celine Yang, Michael A. Taffe

## Abstract

**Background:** Cannabis may reduce the nonmedical use of prescription opioids. Causality of polydrug use is difficult to establish from epidemiological data, and thus controlled laboratory models can test whether cannabinoid co-use with opioids can modulate opioid intake.

**Methods:** Male and female rats were trained to intravenously self-administer (IVSA) oxycodone (0.15 mg/kg/infusion) during 6 h sessions. Separate groups were injected with the vehicle or with THC (5 mg/kg, i.p.; N=10) 30 minutes before sessions for the first three weeks. Treatments were swapped in the fourth week. One male group was trained in the intracranial self-stimulation (ICSS) procedure and assessed for brain reward thresholds prior to each IVSA session.

**Results:** THC treated animals self-administered less oxycodone during acquisition, with a larger differential expressed in the female group. Tolerance to the THC effect developed over the initial weeks, and increasing the dose of THC (10 mg/kg, i.p.) prolonged the suppressing effect on IVSA. While ICSS thresholds increased with sequential IVSA sessions, no differences between THC- and Vehicle-treated groups were observed. Oxycodone IVSA was increased following the first 60 h abstinence interval in THC-treated, but not vehicle-treated, rats. Acute injection of THC, when all animals had been THC abstinent for several weeks, *increased* breakpoints in a Progressive Ratio procedure.

**Conclusion:** These data support the interpretation that THC enhances the reinforcing efficacy of a given dose of oxycodone and may therefore increase the addiction liability.

## Introduction

Overdose fatalities associated with nonmedical use of opioids (Mattson et al., 2021; Spencer et al., 2022) are a small subset of the overall opioid misuse crisis, since these drugs have substantial potential for protracted dependence and use, as reviewed (Leung et al., 2022). Epidemiological data showing a reduction of opioid medical use and/or opioid-related harms in States within the US which have legalized cannabis (Kim et al., 2016; Nguyen et al., 2023; Pardo, 2017; Piper et al., 2017; Shi, 2017) leads to a general hypothesis of the opioid-sparing effects of cannabis (Nielsen et al., 2022). Additional prospective studies likewise show that cannabis can reduce medical use of opioids (Benedict et al., 2022), perhaps unsurprisingly because control of pain is one of the major reasons given for medical cannabis use (Hameed et al., 2023). There are clinical trials registered or ongoing which are designed to determine if medical cannabis can reduce the use of opioids (Jashinski et al., 2022; Zylla et al., 2021), including the specific use of oxycodone in non-cancer pain (van Dam et al., 2023). However, additional epidemiological studies have failed to replicate beneficial findings (DeShea et al., 2023; Kim et al., 2022), highlighting the limitations of epidemiological and even of prospective investigations (Ang et al., 2023; Chihuri and Li, 2019; Vyas et al., 2018). Furthermore, there are often complex outcomes, for example the highest pain relief achieved in chronic pain patients was from combined use of cannabis and opioids compared to either alone, but there was no indication of lower opioid use (Mun et al., 2022). Additional studies report interactions where medical cannabis authorization in pain patients *increased* prescription opioid use in low-intensity users and *decreased* use in those with greater opioid use (Lee et al., 2021). Patients and clinicians express some concerns about cannabis use for pain and the potential for addiction (Cooke et al., 2019), and young adults who use cannabis are at increased risk of nonmedical use of opioids (Rhew et al., 2023). This further emphasizes the need for controlled laboratory studies to more fully determine if cannabis, or its active constituents, can reduce opioid-related harms.

Determination of whether a history of cannabis use leads to increased liability for opioid use or dependency is difficult to determine from human populations, although a recent meta-analysis and systematic review reported increased risk of both opioid use and an opioid use disorder associated with prior cannabis use (Wilson et al., 2022). The theoretical benefits of cannabis co-use for reducing or delaying opioid addiction are relatively unexplored in systematic investigations in human populations. However, evidence from some animal models suggests that the primary psychoactive of cannabis, Δ^9^-tetrahydrocannabinol (**THC**), and some other full-agonist cannabinoid compounds, can reduce the intravenous self-administration (**IVSA**) of heroin (Maguire and France, 2016) or oxycodone (Nguyen et al., 2019). There are competing explanations for why cannabinoid agonists might reduce the number of infusions of an opioid that are obtained in a self-administration model. It may be that the cannabinoid reduces the drive for the opioid, by partially relieving the demand state, or it somehow increases the efficacy of the opioid that is ingested. Evidence from our prior studies tends to support the interpretation that THC enhances the effect of a unit dose of a self-administered opioid. Perhaps most tellingly, acute THC pretreatment *increased* responding under a Progressive Ratio (PR) schedule of reinforcement (Nguyen et al., 2019).

Much of the relevant prior pre-clinical work has been from Maguire, France and colleagues, using a rhesus monkey model of short-access intravenous self-administration of opioids such as remifentanil (Maguire and France, 2020), (Maguire and France, 2018), fentanyl (Carey et al., 2023) or heroin (Maguire and France, 2016; Maguire et al., 2013). Monkeys were assessed in four 25-minute sessions per day in which only 7 infusions were permitted. Our lab has similarly investigated oxycodone IVSA using 1-hour sessions per day (Nguyen et al., 2021). While the short access condition has significant strengths as a model of acute drug reward, it may not best address the ongoing cycle of intoxication / withdrawal that leads to an addictive use disorder (Koob, 2022; Koob and Le Moal, 2001). Extended access to intravenous self-administration of opioids leads to escalating intake and numerous alterations of brain and behavior that are consistent with an opioid use disorder (Ahmed et al., 2000; Blackwood et al., 2019b; Fu et al., 2022; Guha et al., 2022; Kenny et al., 2006; Kimbrough et al., 2020; Matzeu and Martin-Fardon, 2020; Salisbury et al., 2020).

Our model of oxycodone IVSA in daily extended access (“Long Access”; LgA) sessions, scheduled Monday-Friday with weekend breaks, leads to group escalation of intake which includes a marked increase on each Monday over as many as the first three weekends of acquisition. The effect is most pronounced in 11-12 hour daily access models (Nguyen et al., 2021; Nguyen et al., 2017), but it also appears to lesser extent in 8 h (Nguyen et al., 2020a; Nguyen et al., 2019) and even in 1 h (Nguyen et al., 2018) daily access paradigms. Similar patterns have been present in oxycodone IVSA data published by other groups (Blackwood et al., 2019a; Blackwood et al., 2021; de Guglielmo et al., 2020; Illenberger et al., 2023a; Illenberger et al., 2023b; Kallupi et al., 2020; Matzeu and Martin-Fardon, 2020), and while additional studies are insufficiently precise in methods for full clarity, inspection of the acquisition pattern for infusions suggests a similar increase in intake after a weekend abstinence interval (Fu et al., 2022; Kimbrough et al., 2020). Our model also showed that LgA oxycodone IVSA leads to increased brain reward thresholds across a week of sequential sessions of intracranial self-stimulation (ICSS), followed by a return towards baseline conditions after 60 h abstinence (Nguyen et al., 2021). In a critical study, the administration of THC before the IVSA session reduced oxycodone consumption but did not alter the pattern of elevated reward threshold across four sessions (Nguyen et al., 2021). This suggests that THC plus a lower amount of oxycodone produced the same intoxication / withdrawal cycle as a higher amount of self-administered oxycodone without THC. The latter finding was a substitution study, involving only 4 total IVSA sessions with THC, and it was conducted long after the establishment of the stable IVSA and ICSS patterns. Correspondingly, the present study sought to further test the hypothesis that THC enhances the functional efficacy of a given amount of self-administered oxycodone. The design furthers previous work by examining the impact of THC in a between-groups study wherein one group was treated with THC before every self-administration session during acquisition.

## Methods

### Subjects

Subjects used for this study were 40 male and 20 female Wistar rats (Charles River), aged 10-11 weeks on arrival in the laboratory in Cohorts of 20. The vivarium was kept on a 12:12 hour reversed light-dark cycle, and behavior studies were conducted during the vivarium dark period. The animals were pair housed (by treatment condition, see below) and food and water were provided ad libitum in the home cage and in the self-administration chambers. Animal body weights were recorded ∼weekly, beginning at entry to the lab and continuing through the end of the study. Experimental procedures were conducted in accordance with protocols approved by the Institutional Animal Care and Use Committee of the University of California, San Diego and consistent with recommendations in the NIH Guide (Garber et al., 2011).

### Drugs

(-)-Oxycodone HCl was obtained from Sigma-Aldrich (St. Louis, MO) and Δ^9^-tetrahydrocannabinol (THC) was obtained from NIDA Drug Supply. For injection, THC was prepared as a suspension in a 1:1:18 ratio of ethanol:cremophor:saline. The 10 and 20 mg/kg doses were achieved by multiplying the injection volume at the same concentration. Brevital sodium (McKesson Medical-Surgical, Inc. Jacksonville, FL) was injected i.v. at a dose of 3-6 mg/kg to evaluate catheter patency.

### Intracranial Self-Stimulation (ICSS)

The first Cohort (N=20) of male rats was initially prepared for ICSS using surgical preparation, training and maintenance evaluation as previously described (Aarde et al., 2017; Nguyen et al., 2016; Nguyen et al., 2021). In brief, rats were anesthetized with an isoflurane/oxygen vapor mixture (isoflurane 5% induction, 1-3% maintenance) and prepared with unilateral electrodes aimed at the medial forebrain bundle (coordinates: AP -0.5mm, ML ±1.7mm, DV skull -9.5mm). Rats were trained in a procedure adapted from the discrete-trial current-threshold procedure (Kornetsky et al., 1979; Markou and Koob, 1992) until stable ICSS thresholds were exhibited. Thereafter, these rats were surgically implanted with intravenous catheters, see below, and allowed to recover for a minimum of 1 week. ICSS training was resumed for at least one week to re-establish baseline thresholds and thereafter rats were randomly assigned to Vehicle (N=9) and THC (N=10) pre-treatment conditions.

### Intravenous Self-Administration (IVSA)

In brief, rats were prepared with chronic indwelling intravenous catheters using sterile procedures under gas anesthesia as described previously (Nguyen et al., 2017) and in the **Supplementary Methods**. Briefly, the catheter tubing was passed subcutaneously from a port at the back and inserted into the jugular vein. A minimum of 7 days was allowed for surgical recovery prior to starting the experiment. Catheters were flushed with ∼0.2-0.3 ml heparinized (166.7 USP/ml) saline before sessions and ∼0.2-0.3 ml heparinized saline containing cefazolin (100 mg/mL) after sessions. Drug self-administration was conducted in operant boxes (Med Associates) located inside sound-attenuating chambers located in an experimental room (ambient temperature ∼22 ± 1 °C; illuminated by red light) outside of the housing vivarium. To begin a session, the catheter fittings on the animals’ backs were connected to polyethylene tubing contained inside a protective spring suspended into the operant chamber from a liquid swivel attached to a balance arm. Each operant session started with the extension of two retractable levers into the chamber. Following each completion of the response requirement (response ratio), a white stimulus light (located above the reinforced lever) signaled delivery of the reinforcer and remained on during a 20-sec post-infusion timeout, during which responses were recorded but had no scheduled consequences. If no responses were made within 30 min, a single priming infusion was delivered. Drug infusions were delivered via syringe pump. The training dose (0.15 mg/kg/infusion; ∼0.1 ml/infusion) was selected from prior self-administration studies (Nguyen et al., 2018; Nguyen et al., 2017; Wade et al., 2015) and the session duration was 6 hours to match the anticipated maximum duration of THC action (Nguyen et al., 2019).

### Cohort 1

Animals were permitted access to oxycodone IVSA, 0.15 mg/kg/infusion in daily (M-F) sessions of 6 h duration after the ICSS evaluation on each experimental day. Separate groups were injected with the vehicle (N=9) or with THC (5 mg/kg, i.p.; N=10) after the ICSS session and 30 minutes before each self-administration session. In Week 4, the treatment conditions were switched from the Wed session (Session 18) onward and the dose was increased to 10 mg/kg for the Thursday and Friday sessions (Sessions 24 and 25) in Week 5. After a week of no evaluation on ICSS or IVSA, rats were restarted with the same (swapped) pre-treatment. After two days, animals were next evaluated with the 5 mg/kg THC, or Vehicle, administered before the ICSS session in a counterbalanced order; no IVSA sessions were run after these days. These probes followed IVSA sessions 27 and 28. The routine pre-IVSA injections before IVSA sessions were discontinued for Sessions 29-51. Seven days with no IVSA or ICSS sessions elapsed between Sessions 29 and 30 due to unavoidable personnel unavailability. Sessions 32 and 34 were 1 h oxycodone (0.15 mg/kg/infusion) IVSA sessions including either Vehicle or THC injected 30 min prior to the session. Sessions 37 and 39 were 1 h IVSA sessions with only *saline* infusions available, including either Vehicle or THC injected 30 min prior to the session. In all four sessions the ICSS procedure was completed after the IVSA session, and in Sessions 34, 37 and 39 a pre-IVSA ICSS session was also run.

#### Fixed Ratio 5 (FR5)

Further experiments were conducted to probe a persisting small mean difference in IVSA infusions across the original groups that was noticed in Sessions 40-47, conducted without any prior THC or vehicle injection. The response contingency was incremented to FR5 for Sessions 48-56. All subjects were injected with THC (5 mg/kg, i.p.; N=10) 30 minutes before self-administration sessions 52-55.

#### Progressive Ratio (PR)

On Sessions 57-59 animals were tested on a PR procedure with 0.15 mg/kg/infusion oxycodone available. Sessions were a maximum of 3 h in duration. In the PR paradigm, the required response ratio was increased after each reinforcer delivery, within a session (Hodos, 1961; Segal and Mandell, 1974) as determined by the following equation (rounded to the nearest integer): Response Ratio=5e^(injection number\**j*)–5 (Richardson and Roberts, 1996). In this study the *j* value was set to 0.2. Animals were injected with either Veh or THC (5 mg/kg, i.p.) before the 57^th^ and 58^th^ session in a counter-balanced order. The 59^th^ and 60^th^ session were conducted under PR, without any pre-session treatment. On sessions 61-64 animals were tested on a PR procedure with 0.06 mg/kg/infusion oxycodone available. Animals were injected with either Veh or THC (5 mg/kg, i.p.) before the 61st and 62nd sessions in a counter-balanced order. The 63^rd^ and 64^th^ sessions were conducted under PR, without any pre-session treatment. The interval between sessions 59 and 60 was 4 days, following five days of drug access and the interval between the 63^rd^ and 64^th^ was 3 days, following four days of drug access, thus this “weekend” effect was analyzed. Session 65 was a PR session with 0.06 mg/kg/infusion available following a 1-week interruption in running due to a change in vivarium housing equipment.

#### Fixed Ratio 1 (FR1)

The rats continued to self-administer in three-hour sessions under FR1 for Sessions 66-69 with the available oxycodone 0.0., 0.06 or 0.15 mg/kg/infusion in a counterbalanced order. The session duration was selected to maximize comparison of dose sensitivity with the PR procedure.

### Cohort 2

Animals were permitted access to oxycodone IVSA, 0.15 mg/kg/infusion, in daily sessions of 6 h duration. Separate groups were injected with the vehicle (N=9) or with THC (N=9) 30 minutes before each self-administration session. [The catheters were occluded in one animal in each group prior to start of IVSA and catheter repair attempted. Of these, one Vehicle Group individual was started on Day 7 (for the others) and only included for analysis in the subsequent post-acquisition studies. The other repair was unsuccessful and the individual was discontinued from the study.] The first 11 sessions were conducted on *sequential* days, with a two-day break between Sessions 11 and 12, and the THC dose was 5 mg/kg, i.p. for the first 9 sessions. Thereafter the dose was incremented to 10 mg/kg, i.p. for Sessions 10-13 and 20 mg/kg, i.p. for Sessions 14-19. On session 20, the Vehicle group was switched to receive 5 mg/kg THC, i.p., 30 minutes prior to the sessions and the THC group was administered vehicle only. Injections were discontinued on Session 27 and two 6 h sessions without any pre-session injection were conducted. The response contingency was incremented to FR5 for Sessions 29-32

#### Progressive Ratio (PR)

On sessions 33-36 animals were tested on a PR procedure with 0.15 mg/kg/infusion oxycodone available. Sessions were a maximum of 3 h in duration. Animals were injected with either Veh or THC (5 mg/kg, i.p.) before the 33^rd^ and 34^th^ session in a counter-balanced order. The 35^th^ session was conducted under PR, without any pre-session treatment. The per-infusion dose of oxycodone was changed to 0.06 mg/kg/inf for Sessions 36-39 and the rats were injected with either Veh or THC (5 mg/kg, i.p.) before the 36th and 37th sessions in a counter-balanced order. For analysis, the day following the injection days was used as the no-injection comparison.

#### Fixed Ratio 1 (FR1)

The rats continued to self-administer in three-hour sessions with 0.06 mg/kg/inf available under FR1 for Sessions 40-42, and 0.15 mg/kg/inf for Sessions 43-50. Animals were injected with either Veh or THC (5 mg/kg, i.p.), or received no injection, before Sessions 40-42 and again before Sessions 45-47, in a counter-balanced order. Rats were injected with either Veh or THC (10 mg/kg, i.p.) before Sessions 49-50 in a counter-balanced order. To assess the impact of per-infusion oxycodone dose under the FR1 for comparison with the other cohorts, the no-injection days from Sessions 40-42 (0.06 mg/kg/infusion) were compared with Sessions 43 (one day after the mixed-order) and 44 (4 days after the mixed order injections), on both of which 0.15 mg/kg/infusion was available.

### Cohort 3

Female rats were permitted access to oxycodone IVSA, 0.15 mg/kg/infusion in daily sessions of 6 h duration. Separate groups were injected with the vehicle (N=7) or with THC (N=8) 30 minutes before each self-administration session. [Two animals were lost due to mechanical failure of vivarium equipment prior to the start of IVSA training. The three other animals were lost to the study due to surgical / implant complications (cagemate chewed port off, failure to recover over three days, difficulty implanting catheter.)]. The first 12 sessions were conducted on *sequential* days, with a two-day break between Sessions 12 and 13, and the THC dose was 5 mg/kg, i.p. for the first 10 sessions.

Thereafter the dose was incremented to 10 mg/kg, i.p. for Sessions 11-14 and to 20 mg/kg, i.p. for Sessions 15-19. The female rats were assessed for tail-withdrawal latency (from 52°C water) and rectal temperature on the first day after Session 19, with the impact of 10 mg/kg THC, i.p., assessed, and three days later for the impact of 20 mg/kg THC, i.p., in all rats. The measurements were made prior to injection, and 30 and 90 minutes after injection. The 20^th^ IVSA session was initiated the day after this last challenge (six days after Session 19) with no injections delivered prior to the session. For Sessions 22-28, the original Vehicle group was injected with 5 mg/kg, THC, i.p., prior to the IVSA sessions and the original THC group was injected with the vehicle. The response requirement was incremented to FR5 for Sessions 30-32.

#### Progressive Ratio (PR)

On sessions 33-37 animals were tested on the PR procedure with 0.15 mg/kg/infusion oxycodone available. Sessions were a maximum of 3 h in duration. Animals were injected with either Veh or THC (5 mg/kg, i.p.) before the 34^th^ and 35^th^ sessions in a counter-balanced order. The 36^th^ and 37^th^ sessions were conducted under PR, without any pre-session treatment. The per-infusion dose of oxycodone was changed to 0.06 mg/kg/inf for Sessions 38-41 and the rats were injected with either Veh or THC (5 mg/kg, i.p.) before the 39^th^ and 40^th^ sessions in a counter-balanced order. For analysis, the day following the injection days was used as the no-injection comparison.

#### Fixed Ratio 1 (FR1)

The rats continued to self-administer in three-hour sessions under FR1 for Sessions 42-51. Initially, animals had the 0.06 mg/kg/infusion dose available. Thereafter the available oxycodone dose was altered (0.0., 0.06 or 0.15 mg/kg/infusion) in a counterbalanced order in Sessions 43-35.

### Data Analysis

Infusions obtained, the percentage of responses directed at the drug-associated lever, brain reward thresholds and Progressive Ratio data (breakpoint, infusions obtained, percent drug-associated responses and total correct responses) were analyzed by ANOVA, or by mixed-effect models where there were missing values. Within-subjects factors of Session, Dose or Pre-treatment Condition, and a between-subjects factor for treatment Group (THC vs Vehicle) were included where relevant. ICSS data were expressed as a percentage of the individual baselines due to wide individual variation in raw µA values common to this procedure. One rat from each treatment group in Cohort 1 was excluded for the initial 3 weeks of acquisition on grounds of being the only rats with drug-associated lever discrimination to average below 50% across the first three weeks; one was correspondingly excluded from the ICSS data, the other did not have a functional ICSS implant. No rats from Cohort 2 or Cohort 3 met this criterion for exclusion. The data for the excluded rats were returned to their groups in the swap week and thereafter, as their lever discrimination had improved above 50% by that time. One Cohort 1 individual was lost to the study during initial ICSS training due to a health concern of unknown etiology that required euthanasia. Two animals (THC group) were excluded from ICSS analysis due to head implants becoming too damaged to connect the electrodes early in the study. In all analyses, a criterion of P<0.05 was used to infer that a significant difference existed. Any significant main effects were followed with post-hoc analysis using Tukey (multi-level factors), Sidak (two-level factors) or Dunnett (change from first session) correction. All analysis used Prism for Windows (v. 9.5.1; GraphPad Software, Inc, San Diego CA).

## Results

### THC-treated subjects self-administer less oxycodone during acquisition

The initial acquisition of oxycodone self-administration produced similar mean infusions and percent of responses directed to the drug-associated lever across the Cohorts (**Figure 2**); also see **Supplementary Figure S2**. The Cohort 1 animals averaged 78-80% of responses directed to the drug-associated lever for the final three sessions, Cohort 2 animals averaged over 80% of responses directed to the drug-associated lever for the final three sessions and the Cohort 3 animals averaged 76.4-82.0% of responses directed to the drug-associated lever for the final three sessions. In Cohort 1, the six hour access IVSA, run five days per week with 2 days off between weeks, was associated with a pattern of ICSS thresholds that were elevated across consecutive days of IVSA but partially returned towards baseline over the two-day break, consistent with a prior finding with 11 h access oxycodone IVSA (Nguyen et al., 2021).

**Figure 1:**
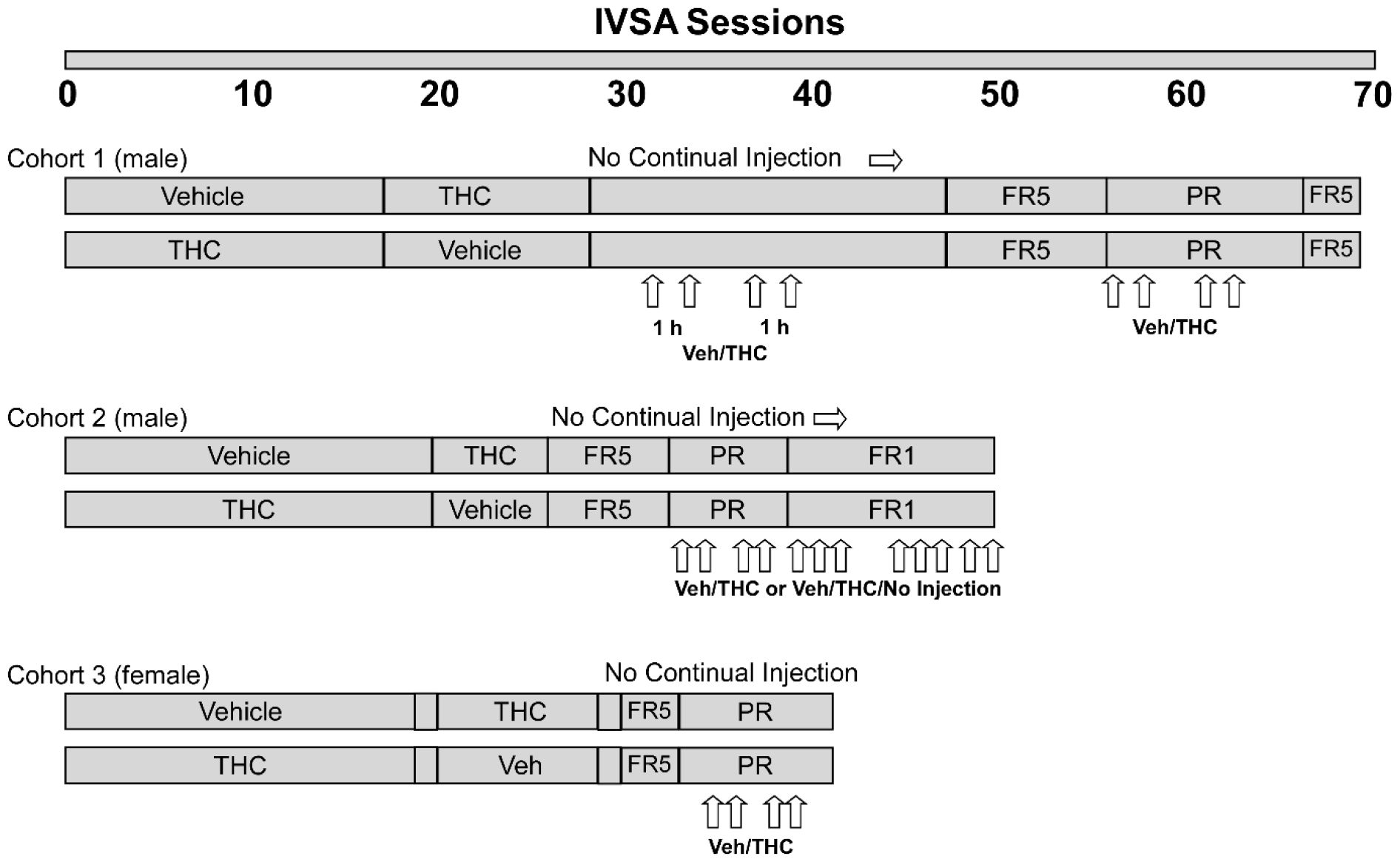
Timeline of experiments for the groups within each Cohort by IVSA session.

**Figure 2:**
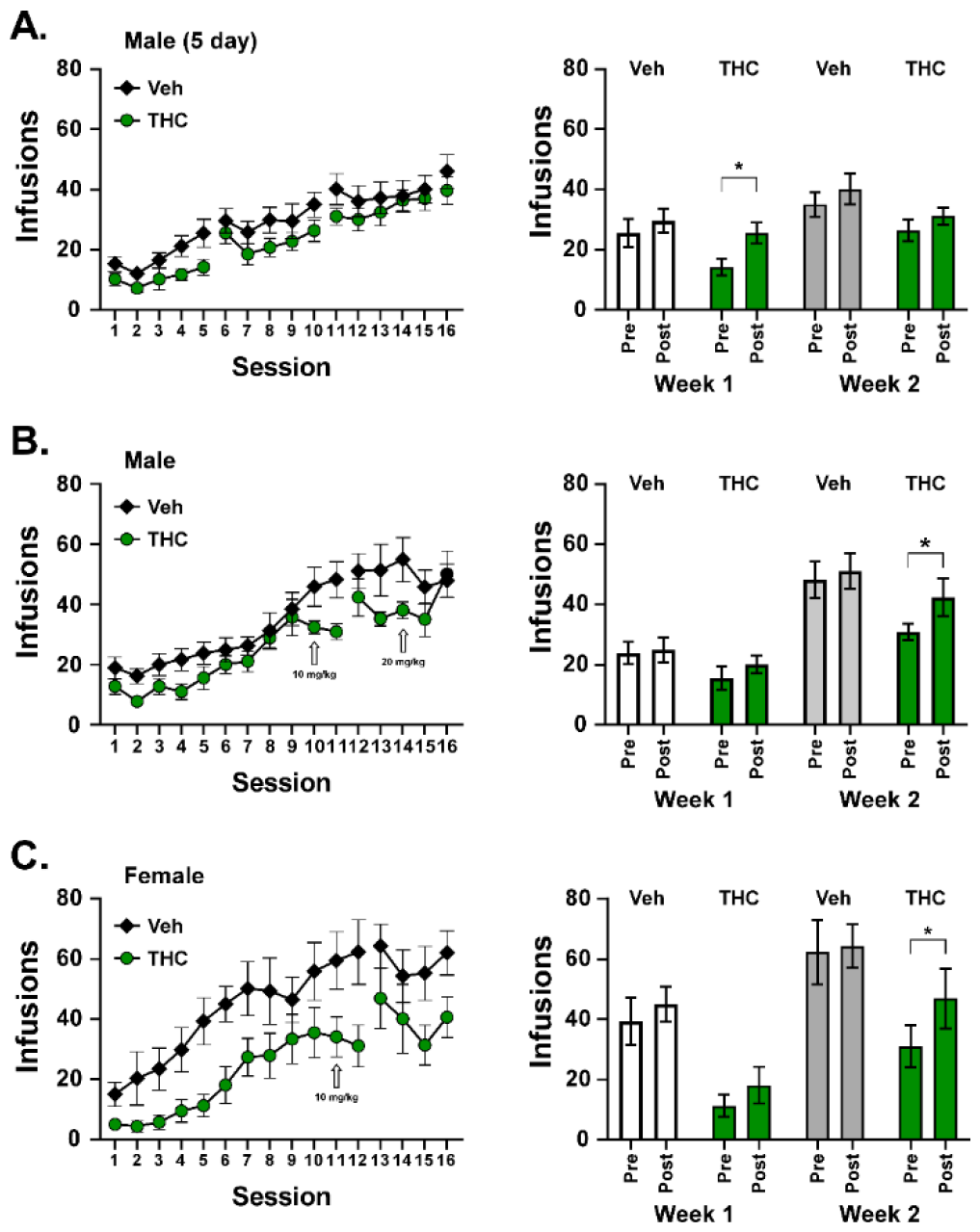
Mean (±SEM) infusions across the first 16 self-administration sessions, and infusions before (Pre) and after (Post) weekend breaks, for groups of A) male rats (N= 17) run 5 days per week and B) male (THC, N=9; Vehicle, N=9) and C) female (THC, N=8; Vehicle, N=7) rats run on sequential days for the first 11 (male) or 12 (female) sessions, followed by a weekend break. The infusions for the weekend break comparison are illustrated in the right panels for <u>Veh</u>icle treated and THC treated subgroups within each cohort. (There was no break between days for Week 1 in panels B and C, see text.) A significant difference from before the weekend break (Pre) is indicated with ^*^.

The data for the 16^th^ session for Cohort 1 is, critically, included on the graph for reference to the effect of weekend breaks, thus the 16^th^ sessions were included for the other cohorts for comparison purposes. Session 16 for Cohort 2 was disrupted because an unscheduled animal facilities water line flush left animals without their usual water source, replaced with gel hydration for two hours prior to the session. This may have caused an increase in self-administration, see **Figure 2B**, consistent with the original intravenous self-administration report showing fluid deprived monkeys would make lever responses for intravenous saline (Clark et al., 1961). Thus, the statistical analysis was limited to 15 sessions for this Cohort.

The two treatment groups within all three cohorts gradually increased the mean number of infusions obtained across the acquisition interval (**Figure 2**), similar to prior results for 8 h, 11 h or 12 h access sessions (Nguyen et al., 2020a; Nguyen et al., 2019; Nguyen et al., 2021). Data were lost, due to computer malfunction, for four animals of Cohort 2 in session 8 therefore all analyses for this cohort that included this session were confirmed by mixed-effects analysis, all others by ANOVA. The analyses confirmed main effects of Session for Cohort 1 [F (15, 225) = 29.75; P<0.0001], Cohort 2 [F (14, 216) = 25.44; P<0.0001] and Cohort 3 [F (15, 195) = 16.64; P<0.0001]. The analysis within cohort confirmed an effect of treatment Group for Cohort 2 [F (1, 16) = 5.12; P<0.05] and for Cohort 3 [F (1, 13) = 6.71; P<0.05] but there were no significant effects of the interaction of factors in any Cohort. The mixed effects analysis of Sessions 1-15 for all male rats (Cohorts 1 and 2) confirmed a significant a significant effect of Group [F (1, 33) = 7.98; P<0.01; Session: F (14, 454) = 47.73; P<0.0001], and the post-hoc test confirmed a difference between the groups only for Session 11. Similarly, when considering all of the male and female rats in the sequential self-administration sessions protocol together (Cohorts 2 and 3), the mixed effects analysis confirmed a significant a significant effect of Group F (1, 31) = 11.06; P<0.005; and of Session: F (14, 426) = 38.36; P<0.0001, but not of the Interaction of Session with Group. The post-hoc test further confirmed specific differences between Groups for Sessions 11 and 12.

For comparison with Cohort 1, the number of infusions obtained by Cohort 2 and Cohort 3 rats are depicted by successive five-day blocks in **Figure 3**, although in this case the first extended break came after Session 11 in the male groups and Session 12 in the female groups. Significant effects of treatment Group were confirmed in the first five sessions for the male [Group, F (1, 16) = 5.59; P<0.05; Session, F (4, 64) = 3.09; P<0.05; Interaction, n.s.] and female [Group F (1, 13) = 7.66; P<0.05; Session, F (4, 52) = 12.29; P<0.0001; Interaction, F (4, 52) = 3.56; P<0.05] rats. Significant effects of Session, but not of Group, were also confirmed for the second five-session segment for female [F (4, 52) = 3.40; P<0.05] and male [F (4, 56) = 18.94; P<0.0001] rats. In the third five-session segment, the analysis confirmed a significant effect of Group for male [F (1, 16) = 5.13; P<0.05] but not for female [F (1, 13) = 4.63; P=0.051] rats.

**Figure 3:**
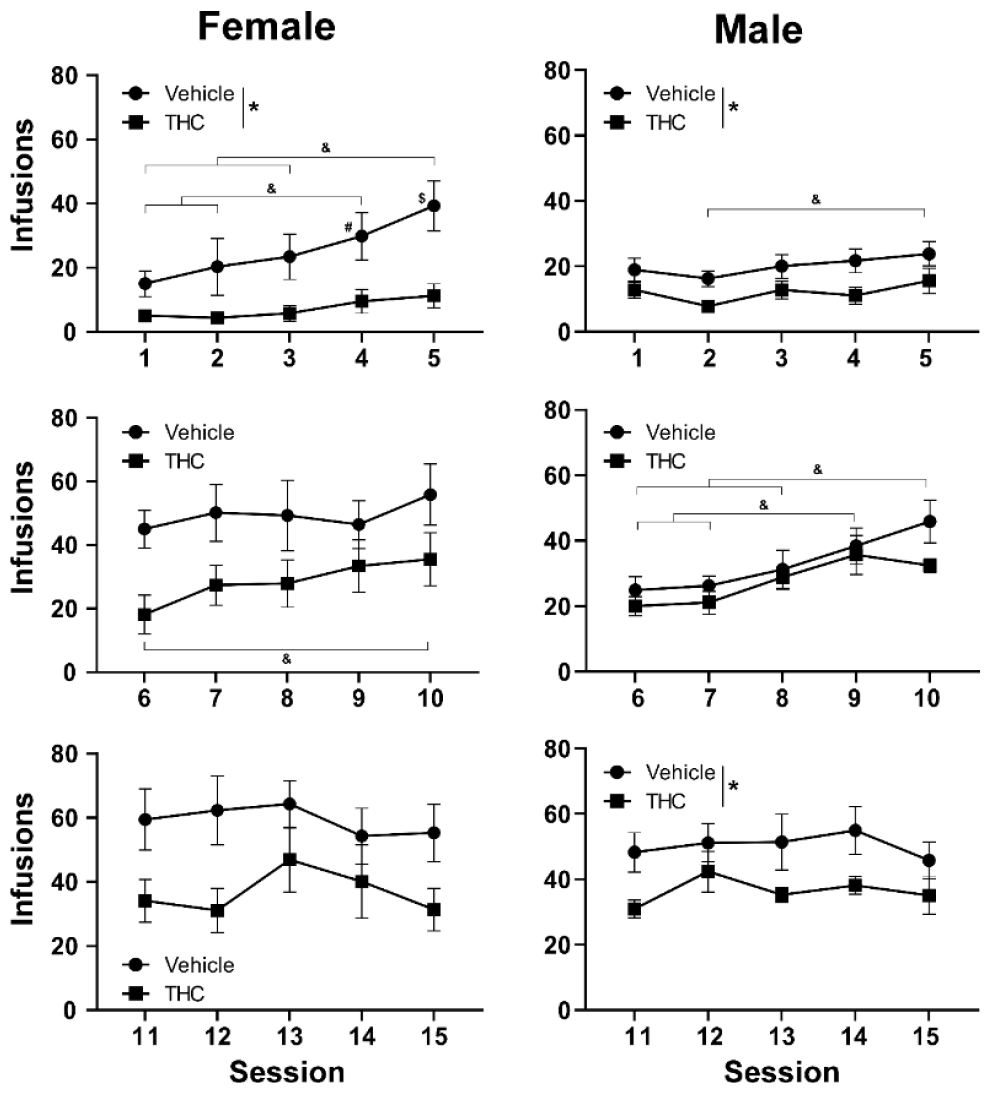
Mean (±SEM) infusions obtained in the first 15 sessions by female and male rats. A significant effect of Group is indicated with ^*^, and a significant effect of Session, collapsed across group is indicated with &.

### Prior THC exposure increases self-administration of oxycodone following weekend abstinence

As in our prior study, the effect of an extended discontinuation over a weekend break was assessed by comparing intake on sessions 5 and 6 for week 1, and on sessions 10/11 (Cohort 1), 11/12 (Cohort 2) or 12/13 (Cohort 3) for week 2.The ANOVA, which included a within subjects factor for Pre/Post break and a between subjects factor for Group/week confirmed significant effects for <u>Cohort 1</u> [Pre/Post: F (1, 30) = 28.63; P<0.0001; Group/Week: F (3, 30) = 3.96; P=0.0171; Interaction: n.s.], and the post-hoc test confirmed there was a significant increase between sessions 5 and 6, but not between sessions 10 and 11. The analysis for <u>Cohort 2</u> also confirmed a significant effect of the weekend discontinuation [Pre/Post: F (1, 32) = 7.85; P<0.01; Group/Week: F (3, 32) = 10.94; P<0.0001; Interaction: n.s.], and the post-hoc test confirmed a significant increase between Sessions 10 and 11, but not Sessions 5 and 6. Finally the analysis for <u>Cohort 3</u> confirmed significant effects [Pre/Post: F (1, 26) = 11.55; P<0.005; Group/Week: F (3, 26) = 7.65; P<0.001; Interaction: n.s.] and the post-hoc test confirmed there was a significant increase between sessions 11 and 12, but not between sessions 5 and 6.

### Prior THC exposure potentiates ICSS reward threshold following daily abstinence in escalated rats

The brain reward thresholds in the Cohort 1 rats increased with each successive day of self-administration within each acquisition week, and the switch week, but there were no differences *between* groups in any of the weeks (**Figure 4**). The analysis confirmed a significant effect of session for ICSS threshold in Week 1 [F (2.71, 37.99) = 4.96; P<0.01], Week 2 [F (2.89, 40.43) = 3.40; P<0.05], Week 3 [F (2.15, 30.08) = 6.14; P<0.01] and Week 4 [F (3.07, 45.98) = 4.10; P<0.05], there was also a significant interaction between Session and treatment Group in Week 2 [F (4, 56) = 2.90; P<0.05]. The analysis confirmed a significant effect of Group on <u>oxycodone infusions</u> in Week 1 [F (1, 15) = 4.97; P<0.05] and a significant effect of the interaction of Group with Session in Week 4 [F (4, 68) = 4.33; P<0.005]. Significant effects of Session on infusions obtained were confirmed in Week 1 [F (2.754, 41.31) = 14.39; P<0.0001] and Week 2 [F (4.015, 60.23) = 14.28; P<0.0001]. The impact of acute THC injection, and self-administration, on ICSS thresholds is described in **Supplementary Materials Figure S7**.

**Figure 4:**
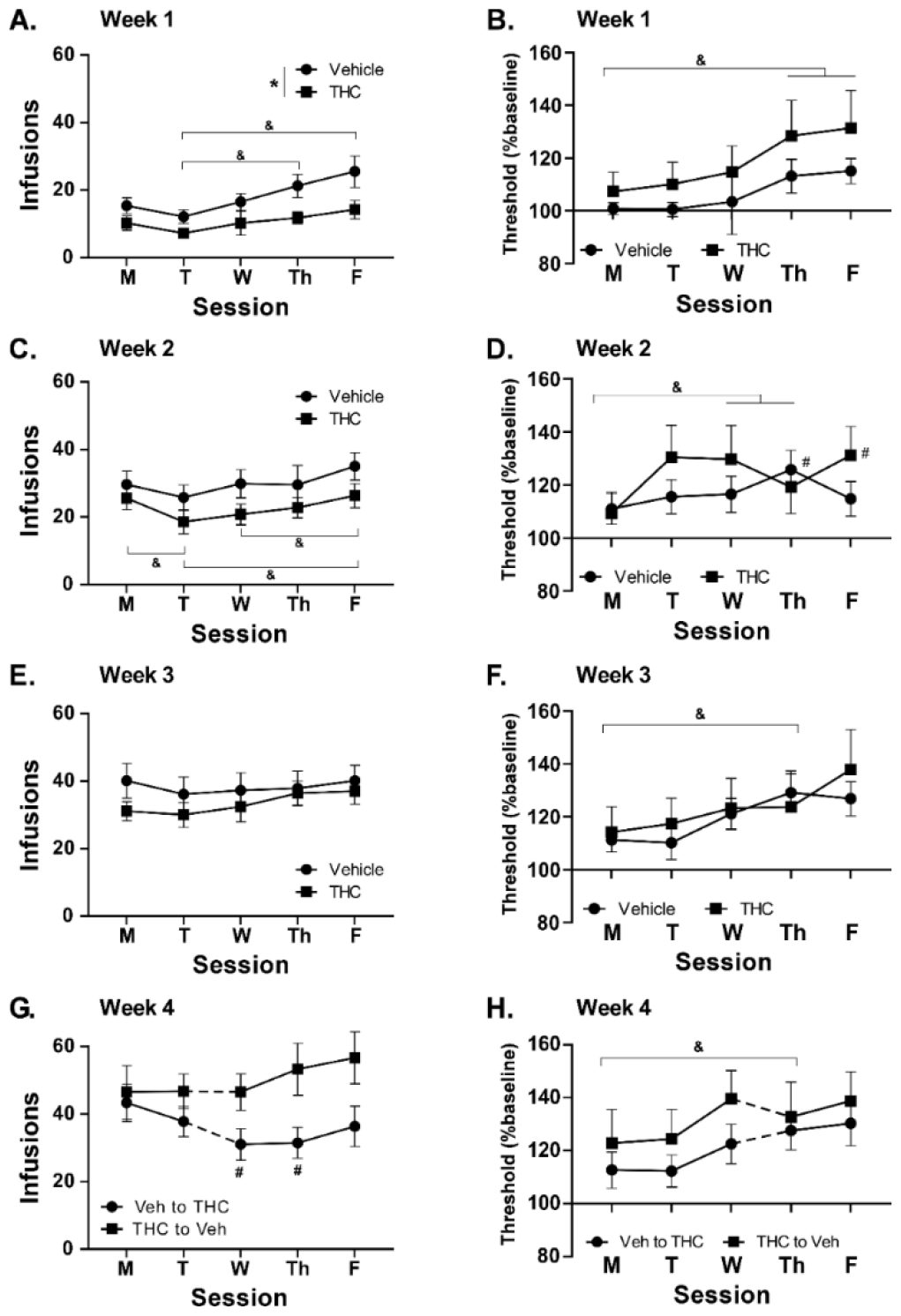
Mean (±SEM) infusions obtained during Weeks 1-4 (A, C, E, G). Mean (N=8 per group Weeks 1-3, N=8 THC to Veh Week 4; ±SEM) ICSS thresholds (as a % of baseline) obtained in Weeks 1-4 (B, D, F, H). The dotted line indicates a switch in the pre-session treatment condition. A significant effect of Group is indicated with ^*^, a significant difference between sessions, collapsed across group, is indicated with &, and a significant difference from Monday, within group with #.

### Relative tolerance to THC may indicate the presence of acute opioid-sparing effects

One possible reason for the more lasting effect of ongoing THC treatment in the female rats is that, contrary to expectation, the THC dosing regimen did not lead to substantial tolerance. To examine this possibility on two measures typically used to assess tolerance, all female rats were challenged with 10 mg/kg THC, i.p., and assessed for rectal temperature and antinociceptive responses (**Figure 5**), one day after the 19^th^ self-administration session. The experiment was repeated four days later with the dose incremented to 20 mg/kg. The THC-treated group exhibited tolerance to the acute temperature disrupting effects, since temperature was only lower than baseline in the Vehicle treated rats, as confirmed by the statistical analysis of the effect of the 10 mg/kg dose [Group: F (1, 13) = 10.61; P<0.01, Time after injection: F (2, 26) = 13.87; P<0.0001, interaction of Group with Time: F (2, 26) = 11.60; P<0.0005] and the 20 mg/kg dose [Group: F (1, 13) = 10.61; P<0.01, Time after injection: F (2, 26) = 13.87; P<0.0001, interaction of Group with Time: F (2, 26) = 11.60; P<0.0005]. The post-hoc test further confirmed that rectal temperature was significantly higher 30 and 90 minutes after injection compared with the pre-injection value in the THC group, and lower 90 minutes after injection compared with the pre-injection and 30-minute post-injection measurements in the Vehicle group. The effects on *<u>nociception</u>* were less clear since the ANOVA confirmed a main effect of Time after injection (F (2, 26) = 12.22; P<0.0005), but not of Group or the interaction of factors. The post-hoc test further confirmed that withdrawal latency was longer 30- and 90-minutes after injection compared with the pre-injection value in the Vehicle group.

**Figure 5:**
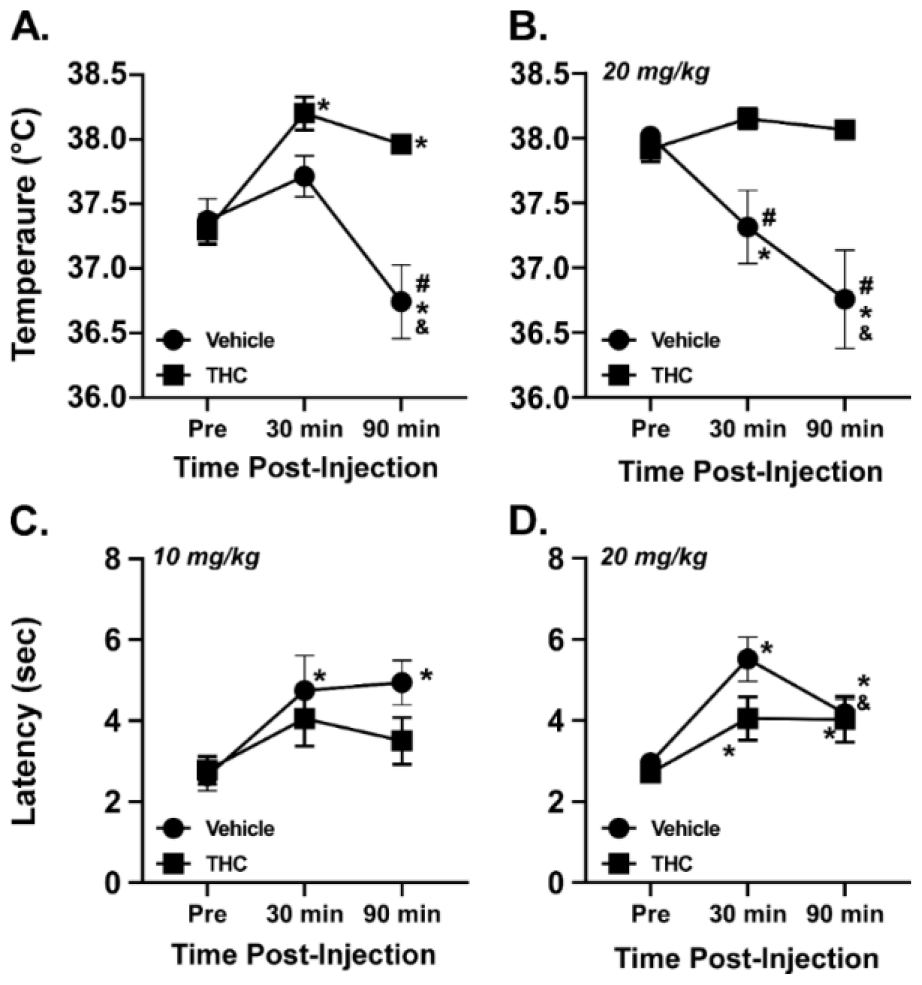
Mean rectal temperature (A., B.) and tail withdrawal latency (C., D.) after injection with 10 mg/kg (A., C.) or 20 mg/kg (B., D.) of THC in groups of rats injected with the Vehicle or THC (5-10 mg/kg, i.p.) before each self-administration session. A significant difference between groups is indicated with #, from the pre-injection value within group with ^*^ and from the 30 minute value within group with &.

#### Progressive Ratio (PR)

The two-way ANOVA, including a factor for the Cohort 1 original groups (N=7 Veh to THC; N=8 THC to Veh), confirmed a significant effect of pre-session Treatment [F (2, 26) = 7.30; P<0.005], but no significant effect of group [F (1, 13) = 1.71; P=0.2140] and no interaction with effect of acute Treatment [F (2, 26) = 0.08; P=0.9250] on breakpoints reached when self-administering oxycodone (0.15 mg/kg/infusion) in the PR procedure. The post-hoc test confirmed that a significantly higher breakpoint was produced after THC compared with either the vehicle injection or the no-injection days, collapsed across group (**Figure 6A**).

**Figure 6:**
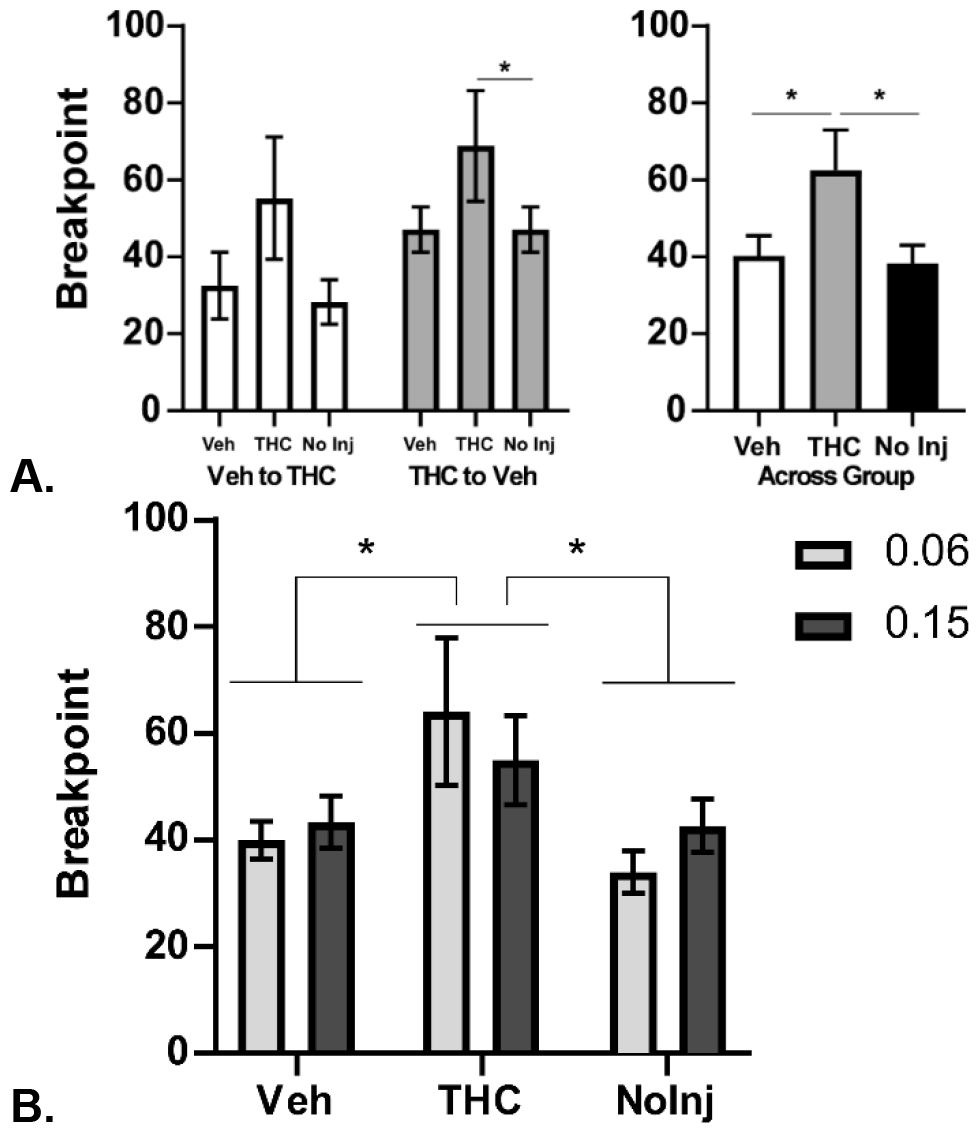
**A)** Mean (±SEM) breakpoint on the PR procedure following injection with Vehicle or THC (5 mg/kg, i.p.) in a randomized order, followed by a no-injection day for the Groups that started with Vehicle injection and were switched to THC (N=7) and the rats that started with THC and were switched to Vehicle (N=8). **B)** Mean (N=25, 12 female / 13 male; ±SEM) breakpoint on the PR procedure following injection with Vehicle or THC (5 mg/kg, i.p.) in a randomized order, followed by a no-injection day. The effect on self-administration of the oxycodone 0.15 mg/kg/infusion dose was evaluated first and the 0.06 mg/kg/infusion dose the following week. A significant difference between pre-treatment conditions, collapsed across oxycodone dose in panel B, is indicated with ^*^.

The impact of THC pre-treatment on the PR procedure was next assessed with all Group 2 and Group 3 animals (N=12 female rats and N=13 male rats). The primary analysis included a factor for pre-treatment condition (Vehicle, THC, no-injection) and another for the available oxycodone dose (**Figure 6B**). One male rat in Cohort 2 did not complete all of the 0.06 mg/kg conditions and thus a mixed effects model was used. The analysis confirmed a significant effect of pre-treatment [F (2, 48) = 7.01; P<0.005] and the post-hoc comparison of the marginal means confirmed significantly higher breakpoints were reached when THC was administered before the session compared with either of the other two pre-treatment conditions.

Follow-up analysis confirmed an overall effect of pre-treatment condition within the female group and within the male group but the post-hoc tests confirmed only a significant difference between THC pre-injection and the no-injection conditions within the male, but not the female, group. Additional follow-up analysis did not confirm any sex differences in breakpoint when either 0.06 or 0.15 mg/kg/infusion was available.

#### Fixed Ratio dose comparison

Altering the per-infusion dose of oxycodone under a FR1 schedule of reinforcement in 3 h sessions confirmed that all Cohorts obtained more infusions of 0.06 mg/kg oxycodone than of 0.15 mg/kg, confirming dose discrimination. See **Supplementary Results Figure S8**.

## Discussion

This study showed first that an acute dose of THC before a 6 h session in which rats are permitted to intravenously self-administer (IVSA) oxycodone reduced the number of infusions obtained. This effect was present in the first week in the male rats in Cohort 1, but was reduced to an unreliable mean difference in Week 2 and disappeared in Week 3 for the chronic administration, between-groups acquisition phase. Introduction of THC to the previously vehicle-treated group of Cohort 1 in Week 4 reduced oxycodone IVSA, which was again attenuated by the following week. Increasing the THC dose from 5 mg/kg to 10 mg/kg appeared to partially sustain the reduction in self-administered oxycodone in Cohort 1. A similar pattern was observed in the male rats in Cohort 2, in this case the mean difference in infusions was maintained by escalating the THC dose from 5 mg/kg to 20 mg/kg across the first 19 sessions. Together this suggested that tolerance to the effect of THC occurs, perhaps after as few as 4 treatment sessions. This interpretation was challenged, however, by the finding in the female THC group of Cohort 3, in which the mean effect of THC to reduce oxycodone self-administration was maintained throughout acquisition. Curiously, physiological tolerance to the temperature reducing effects of THC *was* confirmed in the female rats. In prior work, decrease in oxycodone self-administration was observed in groups of male rats originally trained in 8 h sessions following intermittent THC exposure (Nguyen et al., 2019), as well as in groups trained in 11 h sessions following four sequential days of THC injections (5 mg/kg, i.p.) (Nguyen et al., 2021), albeit long after IVSA acquisition training. THC administered prior to the start of an intravenous self-administration session similarly reduced the amount of oxycodone infusions obtained in groups of male rats trained initially in 8 h sessions (Nguyen et al., 2019) and those trained initially in 11 h sessions, but reduced to 4 h sessions for the THC study (Nguyen et al., 2021). In the former case THC exposure was intermittent, and in the latter case the THC conditions (5 mg/kg, i.p.) were maintained for only four sequential days, however in both of these groups the animals were long past the IVSA acquisition phase. Thus, this current study extends those results to the initial acquisition of self-administration by determining the effect of chronic daily THC across groups.

The male and female rats reached higher breakpoints in the PR procedure after THC (5 mg/kg, i.p.) injection, in a pattern similar to an increased breakpoint reached for heroin IVSA after THC (1 mg/kg, i.p.) in a prior study (Nguyen et al., 2019). The prior intervals of daily THC prior to IVSA sessions produced a lasting tolerance (as demonstrated for Cohort 3 in **Figure 5**), which would explain the different dose (1 vs 5 mg/kg) which resulted in an increase in breakpoints across the two studies. The lack of any effect per-infusion dose of oxycodone in the PR procedure suggests perhaps that in highly trained animals, it is the “hit” associated with each infusion more than the unit dose that is most salient to the animal. However, it was clear in the final FR1 studies in all groups that under this response contingency the animals did discriminate the available per-infusion dose with the expected increase in responding and delivered infusions for the lower dose (**Supplementary Results Figure S8**). Evidence of *increased* behavioral output in a PR procedure, and when the available unit dose per-infusion is lowered in FR (Nguyen et al., 2019), is inconsistent with THC having a non-specific, behaviorally suppressing effect on lever responding. Relatedly, Solinas and colleagues found that heroin IVSA was decreased in a FR procedure while responding was increased in a variable dose procedure after THC (3 mg/kg, i.p.) injection (Solinas et al., 2004). These observations are more consistent with the interpretation that reductions in opioid seeking caused by THC are because THC increases the reward value of a given amount of drug available in each infusion. This interpretation is also consistent with a recent preprint report showing place preference conditioning in mice using combined doses of THC and oxycodone which did not by themselves produce place conditioning (Slivicki et al., 2023). Similarly, this interpretation accords with human laboratory findings in which cannabis smoking increased the abuse-related subjective effects (drug liking, would take again, good drug) of a given dose of oxycodone (Cooper et al., 2018). Finally, this is also consistent with a study reporting that 34.5% of opioid users had ever used cannabis simultaneously with the opioid and that the main reason given was to enhance the feeling of high; only 2.3% reported co-use to diminish withdrawal symptoms (Navaratnam and Foong, 1990).

This study also showed that the impact of THC on IVSA was unlikely to be due to an attenuation of motivational withdrawal. In our prior work we showed that the escalation of oxycodone IVSA in 11-12 hour sessions exhibits an upward ratcheting effect on infusions if a 60 hour discontinuation interval is interspersed with the normative 12 h discontinuation intervals (Nguyen et al., 2021; Nguyen et al., 2017). A more consistent pattern of acquisition was observed in groups run over a similar number of sessions on *sequential* days, i.e., without any 60 hour discontinuation intervals. In addition, treatment with the long-lasting kappa opioid receptor antagonist bridged the 60 hour intervals as if they were merely 12 hour intervals, consistent with the hypothesis that dynorphin/KOR signaling is causally linked to dysphoria and therefore the motivation to IVSA more drug after 60 hours. The present study showed that a similar pattern is produced with 6 h IVSA sessions, followed by ∼18 hours of discontinuation on most days, and a 60 h discontinuation after every 5 sessions, i.e. in Cohort 1. There was no significant increase between sessions 5 and 6 in Cohort 2 and Cohort 3, started only 24 h apart, as with our prior study’s continual 7-day per week access group (Nguyen et al., 2017). However, there *was* a significant IVSA increase in Cohorts 2 and 3 across the first sessions that were separated by a 3-4 day discontinuation interval. This outcome further confirms the specificity of the discontinuation interval in session-to-session increases in IVSA. Of further note in this study was the observation that the effect of the first extended discontinuation from IVSA was largest in the THC-injected groups in all three cohorts. This is consistent with the interpretation that THC increases the efficacy of a unit dose of oxycodone in the sessions leading up to the extended break, thereby creating an increased motivational state after extended abstinence, and generating IVSA levels after the break that are more comparable to levels exhibited by the vehicle-treated groups within the Cohorts that self-administered a higher amount of oxycodone in the prior weeks. N.b. this was the case *despite* the THC groups receiving pre-session THC before the post-weekend sessions. It is likely that this moderated the degree to which IVSA increased after the extended 60 h forced abstinence.

The negative affect theory posits that gradually increasing rates of drug IVSA exhibited with extended daily access sessions arises from dysphoria produced by the cycles of intoxication and withdrawal. This leads to a change in affective setpoint and indeed, foundational work showed that brain reward thresholds measured with the ICSS procedure tend to increase across an interval of extended-access IVSA of several drugs, including heroin (Kenny et al., 2006). The interval of discontinuation is related to the severity of *somatic* signs of withdrawal, which peak across a many day -interval with respect to opioids. This predicts that the escalation of drug self-administration would be affected by the discontinuation interval as well as the intoxication interval, but the former has rarely been directly tested by systematically varying the discontinuation interval. The change in ICSS thresholds observed in Cohort 1 across the acquisition interval was qualitatively similar to the pattern reported previously for 11 h oxycodone self-administering animals (Nguyen et al., 2021). That is, thresholds were gradually elevated across the 5 day week, with repeating cycles of 6 h IVSA followed by ∼18 h discontinuation. Those thresholds then, paradoxically, resolved back towards the baseline (pre-IVSA) values over the 60 hour discontinuation, again similar to a pattern observed in our prior study. Similar to the impact of acute THC treatment (long past the acquisition phase) in the prior study, there was no apparent difference in ICSS patterns associated with the THC-induced reductions in oxycodone infusions across groups in week 1 or in the switch week in Cohort 1 (**Figure 4**). This further supports the interpretation that THC reduces oxycodone IVSA by increasing, rather than decreasing, the subjective value of each infusion.

There was no significant difference between the Cohort 2 and 3 male and female rats in oxycodone IVSA acquisition (**Supplementary Figure S2**), and the slightly poorer discrimination of the males was due to them making more active lever presses during time-out intervals, similar to a prior report (Guha et al., 2022). Increased oxycodone self-administration by female rats has been observed in 8-12 h session IVSA in some prior studies (Kimbrough et al., 2020; Nguyen et al., 2020a), but not in others (Illenberger et al., 2023a). Simple comparisons of infusions acquired under FR1 conditions may be misleading, since introducing a PR schedule eliminates sex differences (Nguyen et al., 2020b), or fails to introduce a sex difference (e.g., Cohort 2 did not differ from Cohort 3 in the PR study). Similarly, increased oxycodone IVSA during acquisition in female rats was accompanied by equivalent scores of somatic withdrawal (Illenberger et al., 2023b). In both cases this suggests an equivalent level of opioid dependence.

In conclusion, this study suggests first that any putative anti-opioid-abuse utility of THC is likely to be transient, because it is highly sensitive to the development of tolerance to THC with regular administration before an opioid self-administration session. This may be a greater likelihood in males than females, although it is unclear how sex differences in rodent responses to THC map on to human sex differences, particularly since the female rats became tolerant to the thermoregulatory and antinociceptive effects of THC. It is possible that escalating the dose of THC would counter this effect, as it did to some degree in Cohort 2 and Cohort 3, but it is unknown how rapidly the dose would have to be increased and for how long dose escalation might be effective in reducing opioid consumption in humans. There was also no evidence in this study that reducing self-administered oxycodone via THC reduces the addiction liability, since similar elevations in reward thresholds after ∼14 hours of drug discontinuation were observed whether oxycodone was self-administered with or without THC injection. Furthermore, THC increased willingness to defend oxycodone intake against increasing workload in the PR procedure and led to increased oxycodone intake after the first extended discontinuation over a weekend in the acquisition phase. Ultimately these data caution against simple interpretations of harm reduction when reduced opioid drug use is associated with cannabis co-consumption.

## Acknowledgements

These studies were supported by the UCSD Center for Medicinal Cannabis Research (Pilot originally awarded to JDN), UCSD Medical Scientist Training Program Summer Undergraduate Research Fellowship (CY), the UCSD PATHS Scholar Program (CY), the Tobacco Related Disease Research Program (T31IP1832; MAT) and the NIH (K99/R00 DA047413; JDN).

## Author contributions

JDN and MAT designed the studies. JDN, YG, and CY performed the research and conducted initial data analysis. JDN and MAT conducted the statistical analysis of the data, created figures, and wrote the paper. All authors approved the submitted version of the manuscript.

## Competing Interests

The authors have no competing interests to disclose.

## Notes

### Competing Interest Statement

The authors have declared no competing interest.

