## Supplementary Materials for "Δ^9^-tetrahydrocannabinol (THC) Increases the Rewarding Value of Oxycodone During Self-Administration in Rats"

Or

Dr. Michael A. Taffe, Department of Psychiatry, 9500 Gilman Drive; University of California, San Diego, La Jolla, CA 92093; USA;

### Supplementary Introduction:

Rats housed in standard laboratory conditions with ad libitum feeding in the home cage continue to gain weight across the adult interval, with male rats growing at a faster rate. Ongoing surveillance of weight gain can serve as a general indication of the impact of treatments, albeit not a very specific indicator, in the absence of close monitoring of food intake.

Rat **oxycodone intravenous self-administration** models have become more plentiful in recent years (Blackwood et al., 2019; Bossert et al., 2018; de Guglielmo et al., 2020; Matzeu and Martin-Fardon, 2020; Mavrikaki et al., 2017; Nguyen et al., 2017; Pravetoni et al., 2014), but the vast majority of knowledge is based on the IVSA of heroin, and to some extent remifentanil. A PubMed search for "rat AND Self-administration AND oxycodone" returned one article from 2005-2009, four articles from 2010-2014, thirty-five articles from 2015-2019 and forty-six articles from 2020 to mid-2023. Given the diversity of daily session intervals deployed in extended access or escalation models in this work (Blackwood et al., 2019; de Guglielmo et al., 2020; Illenberger et al., 2023; Kallupi et al., 2020; Nguyen et al., 2019; Nguyen et al., 2021; Nguyen et al., 2017), it is of significant interest to analyze acquisition parameters for the cohorts as a whole in this six-hour session approach, regardless of treatment group. Similarly, it is of interest to directly compare the sexes, given a diversity of findings for opioid IVSA generally, and for oxycodone IVSA specifically.

**Individual differences** in drug self-administration can potentially provide evidence regarding whether observed effects are related to subjective intoxication and withdrawal states, or are a simple function of the amount of drug exposure, regardless of individually selected satiety point. The THC treatment conditions produced a group mean reduction, however there were individuals from each treatment group in the upper and lower halves of the preference distribution. A median-split analysis was therefore conducted, using the average infusions obtained in the first three weeks for the entire Cohort, regardless of THC treatment.

Sustained elevation of **Intra-Cranial Self Stimulation (ICSS)** reward thresholds has been reported in extended-access paradigms for the self-administration of many drugs including for cocaine (Markou and Koob, 1992), methamphetamine (Jang et al., 2013), nicotine (Kenny and Markou, 2006) and heroin (Kenny et al., 2006). We have shown, however, that in the 5 day on / 2 day off schedule of 11 h oxycodone IVSA, the ICSS pattern is of increasing threshold within week and reset across weekend to almost baseline thresholds (Nguyen et al., 2021). This novel observation is somewhat paradoxical in that the increased self-administration observed after the longer weekend break is hypothesized to result from motivational withdrawal. In our prior study the magnitude of increase in ICSS threshold was related to the IVSA session duration, but this was only determined within group after the 11 h oxycodone acquisition interval. It is therefore of interest to determine if the same general pattern is observed with rats who *initiate* IVSA with 6 h sessions. Evidence also shows that the elevated ICSS thresholds observed prior to the daily ICSS session can be decreased back towards baseline by the self-administration of methamphetamine (Jang et al., 2013) or oxycodone (Nguyen et al., 2021) in long-access trained rats, and by heroin self-administration in 1 h trained rats (Kenny et al., 2006). Thus, a critical test in this study was to determine the impact of a one-hour self-administration session on elevated reward thresholds. This was further complicated by evidence that THC doses of 1 mg/kg or higher, administered acutely by i.p. injection, tend to elevate reward thresholds (Vlachou et al., 2007); a dose of ~ 0.1 mg/kg can reduce thresholds (Gardner et al., 1988; Katsidoni et al., 2013), at least in some laboratories. Since ICSS was assessed before the THC dose during acquisition, it is of further interest to determine if reward thresholds were changed after the 5 mg/kg, i.p., THC dose. This is critical since our prior study found that non-contingent oxycodone did not reduce ICSS thresholds below baseline when administered within a week of prior long access IVSA sessions, but was able to reduce thresholds after 2 weeks of discontinuation from IVSA (Nguyen et al., 2021).

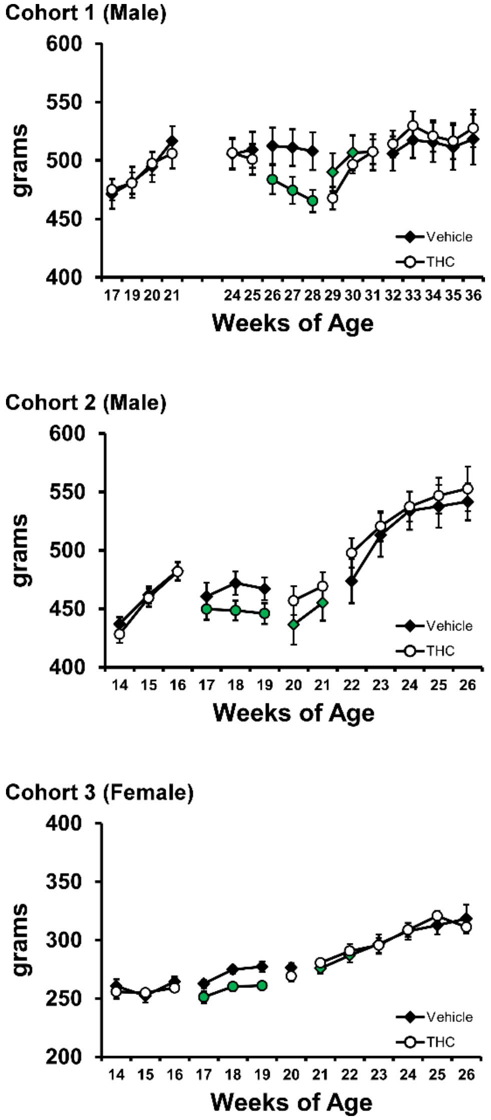


***Figure S1:*** *Mean (±SEM) body weight for the three Cohorts. Critical timepoints include: Cohort 1 (IVSA surgery and recovery- 22-23 weeks; first IVSA day-25 weeks; first day of switched Veh/THC assignment- 28 wks; last day of regular THC/Veh injection- 31 weeks). Cohort 2: (IVSA surgery and recovery- 13-14 weeks; first IVSA day- 16 weeks; first day of switched Veh/THC assignment- 19 weeks; last day of regular THC/Veh injection- 21 weeks). Cohort 3 (IVSA surgery and recovery- 14-15 weeks; first IVSA day- 16 weeks; IVSA suspended for nociception/temperature assessment- 20 weeks; first day of switched Veh/THC assignment- 21 weeks; 22 weeks last day of regular THC/Veh injection).*

### Supplementary Methods:

#### Data Analysis:

A median split analysis was performed for Cohort 1 by ranking individuals, regardless of treatment group, by the average number of oxycodone infusions obtained in the first 15 sessions. This resulted in N=9 Lower Half and N=10 Upper Half animals contributing to the IVSA data. The Lower group included 6 of the THC-treated group, and the Upper Half group included 4 of the THC-treated group. Since 181 was a Lower Half animal and 182 was an Upper animal (both in the THC group but with damage to their ICSS implants) this resulted in N=8 Lower and N=9 Upper Half individuals grouped for the ICSS median split analysis.

In all analyses, a criterion of $P<0.05$ was used to infer that a significant difference existed. Any significant main effects were followed with post-hoc analysis using Tukey (multi-level factors), Sidak (two-level factors) or Dunnett correction. All analysis used Prism for Windows (v. 9.5.1-10.1.0; GraphPad Software, Inc, San Diego CA).

### **Supplementary Results**:

#### THC Decreases Body Weight:

In this study, the administration of THC prior to oxycodone IVSA reduced bodyweight relative to the vehicle-treated group in all three cohorts (**Figure S1**). Weights were obtained on Wednesdays and thus the timing relative to experimental manipulations is approximate. However, since the THC / Vehicle treatment Group swaps were conducted mid-week, the first post-swap weight came after a week of this change. Weight differences were most pronounced in the initial three weeks of THC treatment, but also appeared in the THC/Vehicle swap weeks for the males (Cohort 1 and 2). The female cohort experienced a ~week discontinuation of IVSA prior to the swap (at 20 weeks of age) to facilitate the tolerance study, which may explain the difference. Upon discontinuation from chronic THC treatment before IVSA sessions, all three of the THC-treated groups regained a mean weight nearly identical to the Vehicle-treated group in their Cohort within a week or two.

#### THC Affects Responsivity During Acquisition:

THC produced a small *qualitative* difference in the early stages of acquisition, as more priming infusions (delivered once per session to any individuals for whom the initial 30 minutes of a session elapsed without a lever press). In Cohort 1 six to eight rats received priming infusions in sessions 1-3 (17 total primes to THC group, 3 to the Veh group), two and three rats in sessions 4 and 5, respectively, and a single priming infusion was delivered, across all rats, for sessions 6-17. In Cohort 2 five to ten rats received priming infusions in each of sessions 1-3 (17 total primes to THC group, 5 to the Veh group), two to three rats in each of sessions 4-7 (all THC group), zero primes in Sessions 8-14, and one THC rat on Session 15. In Cohort 3 six to eight rats received priming infusions in each of sessions 1-3 (16 total primes to THC group, 4 to the Veh group), two to five rats in each of sessions 4-7 (1 to a Veh rat, 14 to THC group rats in total), and six total primes in Sessions 8-16 (all THC group rats).

### Acquisition By Cohort:

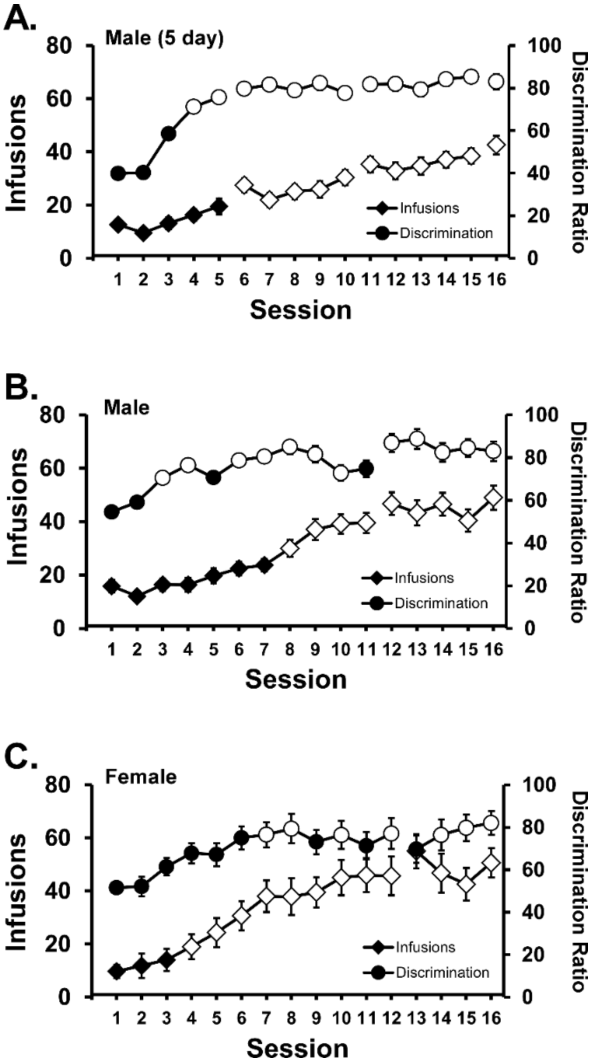


***Figure S2:*** *Mean (±SEM) infusions and discrimination ratio (percent of all responses directed at the drug-associated lever) for all rats in A) Cohort 1 (N=17), B) Cohort 2 (N= 18) and C) Cohort 3 (N= 15). Open symbols indicate a significant difference from the first session.*

Oxycodone infusions obtained and the percentage of all responses directed at the drug-associated lever increased across the initial 16 sessions in all three Cohorts as is depicted in **Figure S2**. Correspondingly, a one-way analysis for all subjects in each cohort confirmed a significant effect of Session on infusions for Cohort 1 [F (15, 240) = 30.97; P<0.0001], Cohort 2 [F (15, 247) = 24.82; P<0.0001], Cohort 3 [F (15, 210) = 17.02; P<0.0001] and a significant effect of Session on percent drug-associated responses for Cohort 1 [F (15, 240) = 14.91; P<0.0001], Cohort 2 [F (15, 247) = 6.30; P<0.0001] and Cohort 3 [F (15, 210) = 4.06; P<0.0001].

### Acquisition Across Cohort:

The three Cohorts differed from each other in either sex or the presence/absence of a 60 h discontinuation after the first five sessions, and therefore it was of interest to determine if there were any substantive differences between Cohorts in terms of overall acquisition. The two-way ANOVAs including factors for Cohort and for Session confirmed that the initial acquisition of oxycodone self-administration produced similar mean infusions and percent of responses directed to the drug-associated lever across the Cohorts, **Figure S3**. In the mixed effect analysis by Cohort, for infusions (**Figure S3A**) there was a significant main effect of Session [F (15, 697) = 63.02;  P<0.0001], and interaction of Session with Cohort  [F (30, 697) = 2.106;  P<0.001]. The post hoc test confirmed that Cohorts 1 and 2 differed only on session 12. Cohort 3 differed from Cohort 1 on Sessions 7, 10, 13 and from Cohort 2 on Session 7. Within Cohort, infusions were significantly higher compared with the first session for Cohort 1 (sessions 6, 8-16), Cohort 2 (sessions 8-16) and Cohort 3 (Sessions 5-16). Tuning of behavior to direct most of the responses to the drug-associated manipulandum (e.g., lever or nosepoke) is often used as evidence of successful acquisition of self-administration. Although such criteria vary across studies, a 2:1 ratio or 80% are two frequently used thresholds. The Cohort 1 animals averaged 78-80% of responses directed to the drug-associated lever for the final three sessions, the Cohort 2 animals averaged over 80% of responses directed to the drug-associated lever for the final three sessions and the Cohort 3 rats averaged 76, 80 and 82% of responses directed to the drug-associated lever for the final three sessions (**Figure S3B**). There was no significant effect of Cohort or of the interaction of Cohort with Session. Post-hoc exploration of the main effect of Session [F (15, 697) = 21.20; P<0.0001] confirmed significantly higher discrimination relative to Session 1 (Sessions 3-16), Session 2 (Sessions 3-16), Session 3 (Sessions 6-16), Session 4 (Session 15) and Session 5 (Sessions 15-16).

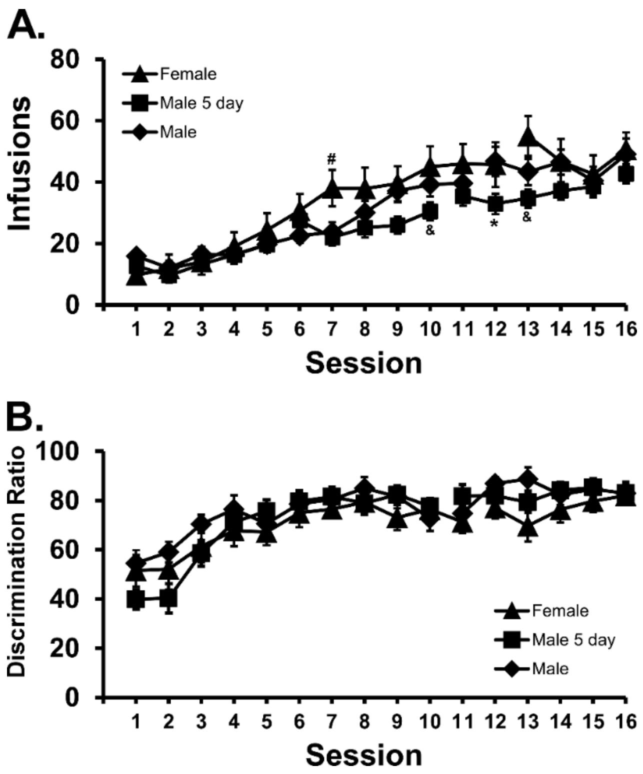


***Figure S3:*** *Mean A) infusions and B) percentage of drug-associated responses for groups of male rats (N= 17) run 5 days per week and male (N=18) and female (N=15) rats run on sequential days for the first 11 (male) or 12 (female) sessions. A significant difference from both male groups is indicated with #, from the female group with & and a difference between male groups with *.*

### Effect of Oxycodone IVSA on ICSS Thresholds in Cohort 1:

In this study, the six-hour access oxycodone IVSA, run five days per week with 2 days off between weeks in Cohort 1, was associated with a pattern of ICSS thresholds that were elevated with consecutive days of IVSA but partially re-set over the two day break (**Figure S4**). The one-way ANOVA confirmed a significant effect of session [F (6.713, 100.7) = 4.27; P<0.0005] on threshold, and the

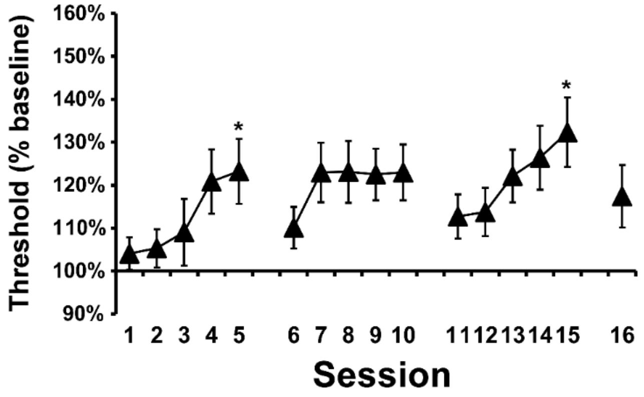


***Figure S4:*** *Analysis of mean (N=17; ±SEM) ICSS thresholds for Cohort 1, collapsed across treatment group, confirms that 6 h IVSA sessions result in gradually increasing threshold across sequential days as previously reported for 11 h IVSA sessions. Data are represented as a percent of an individual 2 day baseline determined post-surgery and before starting IVSA. Two individuals (182/186) were excluded from this analysis since they were excluded from the IVSA acquisition analysis for exhibiting active-lever discrimination of below 50% in first three weeks.*

Dunnett post-hoc test confirmed significant elevations on Session 5 and 15 over Session 1.

**Median Split Analysis (Cohort 1):**

The median split resulted in N=9 Lower-Half individuals and N=10 Upper Half individuals (no animals were excluded for this analysis) for the IVSA data, within which three Vehicle-treated individuals were in the lower half, and four THC treated individuals were in the upper half, of the distribution. One animal in each half of the preference distribution did not have ICSS data due to implant problems, thus N=8 (Lower-Half) and N=9 (Upper-Half) for that analysis. The analysis was conducted for the first three acquisition weeks, the switch week 4 (Groups switched the Vehicle / THC (5 mg/kg i.p.) on Wednesday), and week 5 for which the Tuesday and Wednesday session data were omitted, due to computer failures for half the animals. The THC dose was increased to 10 mg/kg on Th and Fr sessions of Week 5 (**Figure S5**). Analysis by ANOVA confirmed significant effects of Session [F (22, 374) = 26.76; P<0.0001] and of median split Group [F (1, 17) = 13.95; P<0.005] on infusions obtained (**Figure S5A**). Post-hoc analysis further confirmed significant differences between the Groups in sessions 10-13, 16, 24-25. There was a significant effect of Session on the <u>percent of drug-associated responses</u> [F (22, 374) = 7.76; P<0.0001] and on <u>ICSS thresholds</u> [F (24, 360) = 4.4; P<0.0001], but no main or interacting effects of median-split Group were confirmed in the ANOVA for those measures (**Figure S5B, C**).

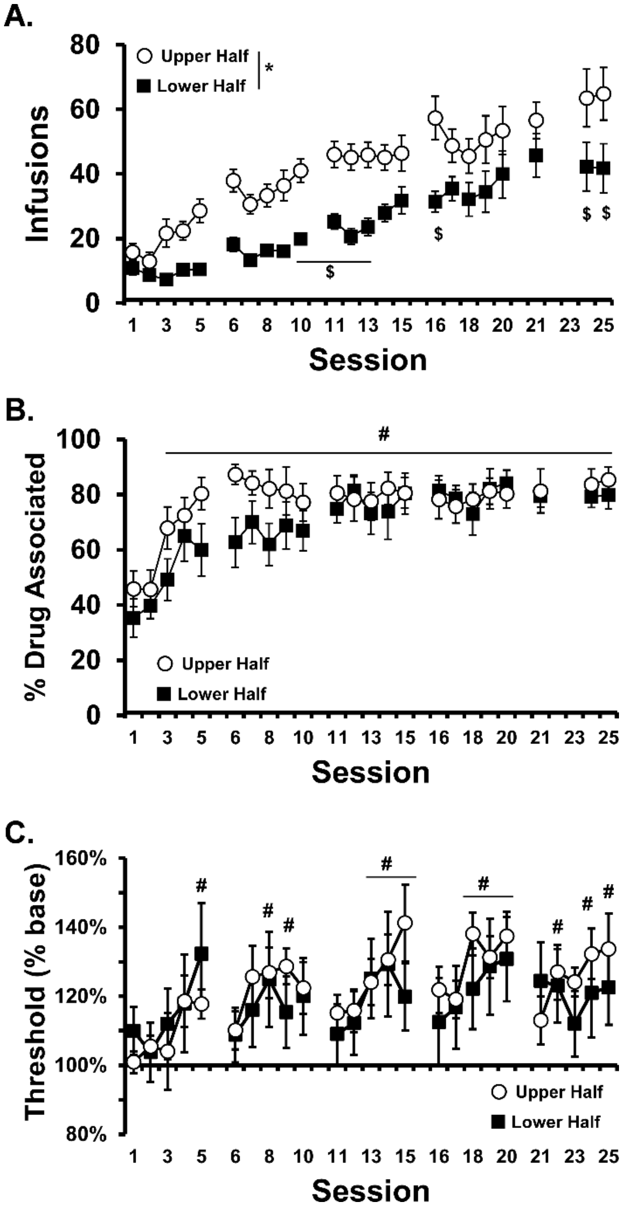


***Figure S5:*** *Mean (±SEM) A) infusions obtained, B) percent responses on the drug-associated lever by the Lower (N=9) and Upper (N=10) IVSA preference subsets of Cohort 1. A significant main effect of Group is indicated with *, a significant difference between Groups for a given session with $, and a significant difference from Session 1, across groups, is indicated with #. C) Mean (±SEM) ICSS thresholds (as a % of baseline) obtained in Weeks 1-6 by Lower (N=8) and Upper (N=9) IVSA preference subsets of Cohort 1. A significant difference from Session 1, across groups, is indicated with #.*

**THC Swap Cohort 2 and 3:**

There was no significant effect of the switch of pre-session treatment (THC vs Vehicle) in Cohort 2 (not shown) or 3 (**Figure S6**). During the injection swapped conditions under the FR1 response contingency there was no change across sessions for THC to Veh or Veh to THC treatment groups within each Cohort, and no significant difference between the treatment groups (**Figure S6**).

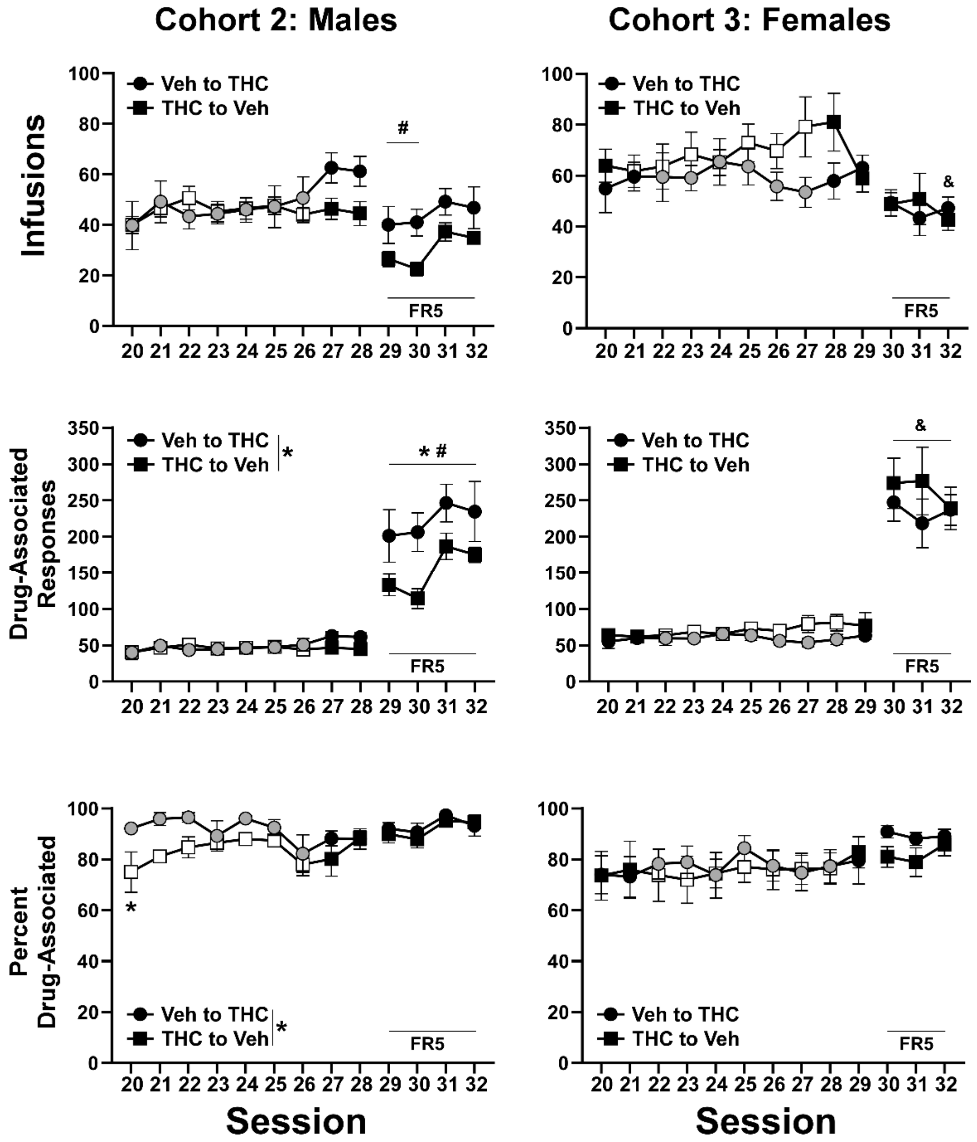


***Figure S6:*** *Mean (±SEM) Infusions, total responses on the drug-associated lever and percent of all responses directed at the drug-associated lever for Cohorts 2 and 3 in 6 h sessions. Sessions preceded by a vehicle injection are indicated with an open symbol and those preceded by a THC injection are indicated with greyed symbols. In Cohort 2, the Veh to THC group was N=5 and the THC to Veh group was N=8. In Cohort 3, the Veh to THC group was N=7 except for the final session which was N=6. The Veh to THC group was N=7 in total with data missing for Sessions 29-32 from 1 subject (found dead of unknown causes) and for Sessions 24-28 for another subject (suspended temporarily due to self-injurious behavior). A significant difference between groups is indicated with *, and a significant difference from the FR1 session immediately preceding the FR5 swap within group with # and across groups with &.*

**Effect of FR1 to FR5 in Cohorts 2 and 3:**

For Cohort 2, the impact of changing from a FR1 to a FR5 schedule of reinforcement differed between the groups (**Figure S6**), despite discontinuation of the pre-session THC. The mixed-effects analyses confirmed significant effects of Session for infusions [F (12, 132) = 6.39, P<0.0001], drug-associated responses [F (12, 132) = 72.10, P<0.0001] and percent drug-associated responses [F (12, 132) = 2.54, P<0.005]. There was also a significant impact of Group on drug- associated responses [F (1, 11) = 5.47, P<0.0001] and percent drug-associated responses [F (1, 11) = 5.49, P<0.05] and a significant effect of the interaction of Session with Group on infusions [F (12, 132) = 2.55, P<0.005] and total drug-associated responses [F (12, 132) = 4.270, P<0.0001]. The original Vehicle group, that had most recently experienced THC prior to IVSA sessions, emitted more drug-associated responses than the original THC group, under FR 5.

For Cohort 3, the mixed-effects analyses confirmed significant effects of session for infusions [F (12, 134) = 4.19, P<0.0001], Drug-associated responses [F (12, 134) = 68.97, P<0.0001] and percent drug-associated responses [F (12, 134) = 2.10, P<0.05]. There were no significant effects of Group or of the interaction of Session with Group for this cohort.

**ICSS: Effects of acute THC treatment and 1 h IVSA of oxycodone**

This experiment in Cohort 1 first tested the effect of Vehicle or THC (5 mg/kg, i.p.) administered *before* the ICSS session on a Wed and a Fri in a counterbalanced order. This was assessed following one week of abstinence from ICSS and IVSA sessions inserted after the 10 mg/kg test of Week 5. During this experiment the rats were maintained in the swapped 5 mg/kg THC (N=9) / Veh (N=8) injection conditions, administered after the ICSS session and before a 6 h IVSA session for the other days of the week (M, T, Th). For the two ICSS test days, there were no IVSA sessions run after the ICSS session. ICSS was evaluated once per day and thus the Pretreatment value was derived from the day immediately prior to the relevant injection day. One individual (176) was not included because it had been reduced from standard 6 h IVSA in this interval due to self-injurious behavior.

The initial three-way ANOVA confirmed a significant effect of Veh vs THC acute injection [F (1, 14) = 25.87; P<0.0005], of Pre/Treatment Day [F (1, 14) = 31.64; P<0.0001] and an interaction of those factors [F (1, 14) = 26.88; P<0.0005] but did not confirm any significant effect of Group or any interaction of Group with other factors on the ICSS thresholds (**Figure S7, left panel**). Collapsed across Group, the follow-up two-way ANOVA again confirmed a significant effect of Veh vs THC acute injection [F (1, 15) = 24.75; P<0.0005], of Pre/Treatment Day [F (1, 15) = 31.74; P<0.0001] and an interaction of those factors [F (1, 15) = 29.41; P<0.0001]. The Tukey post hoc test confirmed that thresholds were significantly elevated after THC injection compared with either the comparison day before or with the vehicle injection day.

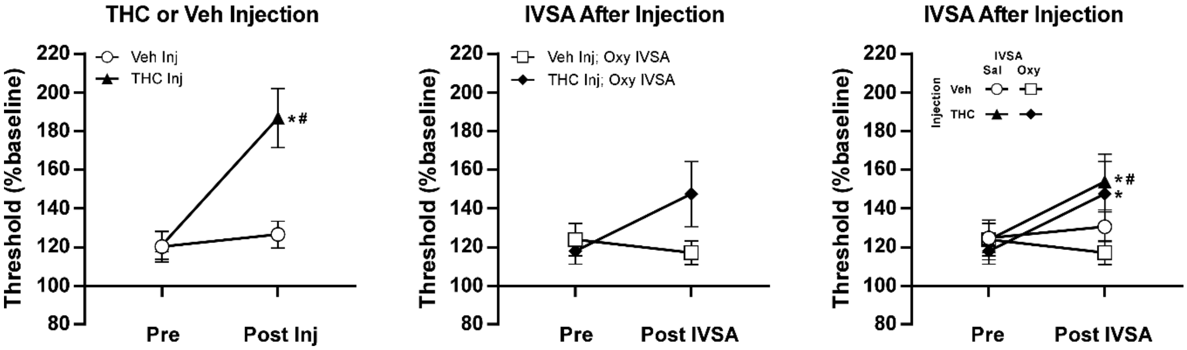


***Figure S7:*** *Mean (±SEM) ICSS thresholds left) before and after injection with THC or the Vehicle; middle) before and after 1 h oxycodone IVSA preceded by THC or Vehicle injection; and right) before and after 1 h oxycodone or Saline IVSA preceded by THC or Vehicle injection. A significant difference from the Pre-injection value is indicated with * and a difference between post-injection THC versus Vehicle injection values with #.*

We next conducted an experiment in which animals completed an ICSS session, and then received an injection of vehicle or 5 mg/kg THC 30 minutes prior to a ***1 h*** oxycodone IVSA session. This was followed with a post-IVSA ICSS session to determine the impact of one hour of oxycodone IVSA on the threshold (**Figure S7, middle panel**). The pre-IVSA injection order was balanced across two sessions run on a Wed and a Fri. During this week, the M, T and Thu days were standard, i.e., ICSS followed by the 6 h IVSA session, in this case with no injection prior to IVSA. For this, and the following, study data were available for N=14 individuals for ICSS. Analysis of the ICSS data compared the Pre and Post IVSA measures across the vehicle and THC injection conditions and the ANOVA confirmed a significant interaction of ICSS session with pre-IVSA injection condition [ F (1, 13) = 5.23; P<0.05]. The post-hoc test did not confirm a significant change in reward thresholds after THC injection relative to the Pre on that day (P=0.86) or relative to the Post-IVSA after the Vehicle injection (P=0.076).

The final experiment was similar, except in this case the self-administration of oxycodone was contrasted with self-administration of saline, with prior injection of either THC or the Vehicle (**Figure S7, right panel**). The three-factor ANOVA confirmed significant effects of Pre/Post [F (1, 13) = 9.54; P<0.01], of THC/Veh injection condition [F (1, 13) = 4.96; P<0.05], and of the interaction of these two factors [F (1, 13) = 8.26; P<0.05]. There were no significant effects of self-administration condition alone or in interaction with other factors in the analysis. The Tukey post-hoc test confirmed that thresholds were significantly higher post-IVSA in the THC injection conditions relative to the respective pre-IVSA measurement. Thresholds were also higher after THC injection and saline IVSA compared with Vehicle injection and either saline or oxycodone IVSA.

### FR1 oxycodone dose-substitution

This study was conducted in all three Cohorts to further determine the sensitivity to oxycodone dose substitution under the FR1 schedule of reinforcement, given there was no apparent difference in responding for the different per-infusion doses under PR. Note that we reported a similar insensitivity of the PR measures to per-infusion dose in a prior study of the effect of THC pretreatment on heroin IVSA (Nguyen et al., 2019). In all three Cohorts it was verified that significantly more infusions were obtained when the per-infusion oxycodone dose was 0.06 mg/kg versus 0.15 mg/kg (**Figure S8**). While the same number of infusions of vehicle were obtained as when 0.06 mg/kg/infusion was available in Cohort 1 and 3, the further increase in time-out responding (on the drug-associated lever) when only vehicle was available indicates behavioral discrimination of the conditions. In Cohort 1, the one-way ANOVAs confirmed a significant effect of per-infusion condition on infusions [F (2, 26) = 12.12; P<0.0005] and Time-Out Correct responses [F (2, 26) = 8.03; P<0.005], but not on Time-Out Incorrect responses, nor on the Percent Drug-Associated responses. [N.b. One animal in Cohort 1 (181) disconnected on first day (0.06), was rerun after the last scheduled day.] In Cohort 2, the one-way ANOVAs confirmed a significant effect of per-infusion condition on infusions [F

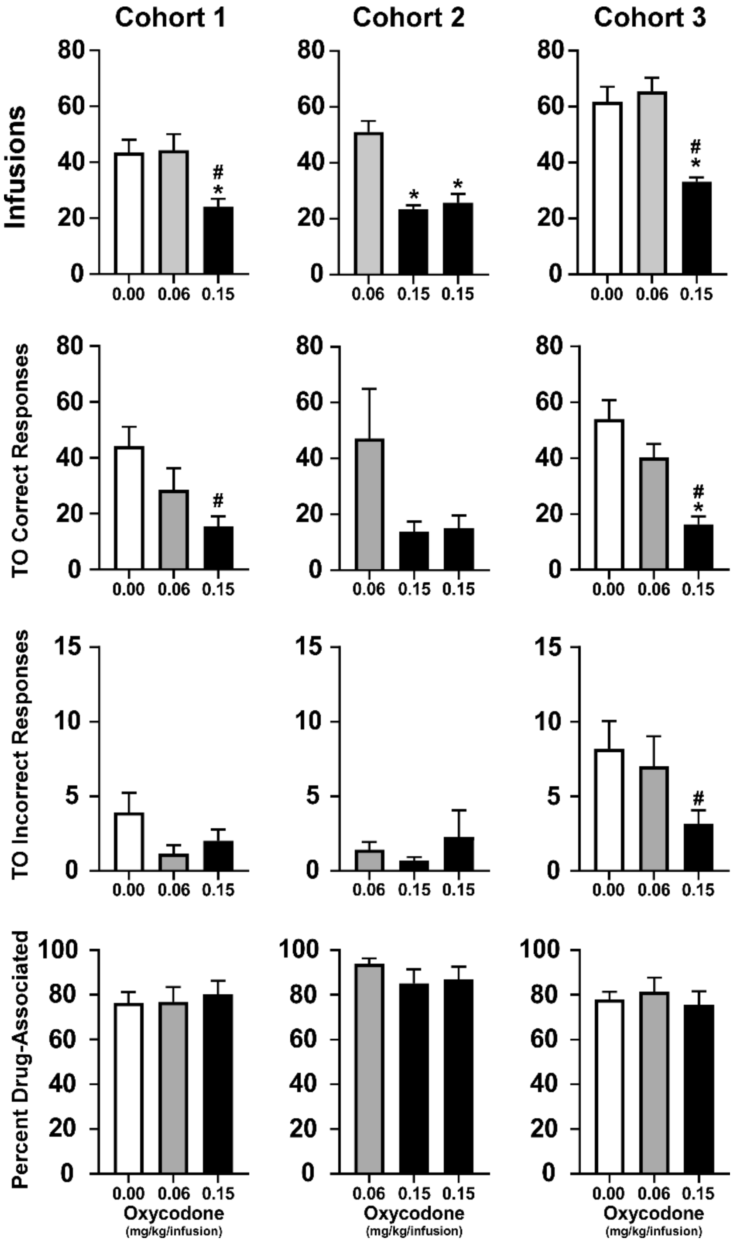


*Figure S8: Mean (±SEM) Infusions, Time-Out Correct Responses, Time-Out Incorrect and Percent Responses on the Drug-Associated lever for Cohort 1 (N=14), Cohort 2 (N=12) and Cohort 3 (N=12) in 3 h sessions under a Fixed-Ratio 1 (FR1) contingency. A significant difference from the 0.06 mg/kg/infusion condition is indicated with * and from the saline (0.00 mg/kg/infusion) condition with #. Correct indicates responses on the drug-associated lever and Incorrect responses on the alternate lever.*

(2, 22) = 58.68; P<0.0001], but not on Time-Out Correct or Time-Out Incorrect responses, nor on the Percent Drug-Associated responses. For this analysis, the 0.06 mg/kg/infusion condition was the no-injection days on the randomized No Injection, Vehicle, THC (5 mg/kg) sequence. The first 0.15 condition was from the day after this sequence. To determine any possible carryover effects of the THC administered on the two prior days, another day of 0.15 mg/kg/infusion conducted three days later (after a weekend break) was included. The analysis for Cohort 3 confirmed a significant effect of per-infusion condition on infusions [F (2, 22) = 19.35; P<0.0001], Time-Out Correct [F (2, 22) = 21.56; P<0.0001] and Time-Out Incorrect [F (2, 22) = 3.92; P<0.05] responses, but not on Percent Drug-Associated lever responses. The Cohort 3 data were previously published in pre-print (Nguyen et al., 2023) and are included here for comparison purposes.

### Supplementary Discussion:

These additional results and analyses further elaborate the impact of the approaches used in this study. All three Cohorts self-administered approximately equal amounts of oxycodone during acquisition, increased their intake on a group mean basis, and refined their lever pressing discrimination to a similar degree and across a similar time scale (**Figure S2**). Repeated THC injection before IVSA sessions produced a small group difference in bodyweight but this was eliminated within a week or two of discontinuing repeated THC (**Figure S1**). All Cohorts were sensitive to the available per-infusion dose when it was altered under a FR 1 contingency (**Figure S8**) and the Cohorts tested on FR 5 schedule of reinforcement significantly increased their responding (**Figure S6**). These adjustments show the groups were sensitive to dose and work requirement, even though a reduction in the per-infusion dose of oxycodone did not significantly alter breakpoints in all three Cohorts in the PR procedure (described in the main report). This was not too surprising since the control group self-administered about 9 infusions at the 0.15 mg/kg dose and 7 at the 0.06 mg/kg dose using the same PR schedule used here in a prior study (Nguyen et al., 2018). Similarly, we also reported that per-infusion doses of 0.06 vs 0.006 mg/kg of heroin produced differential responding in a FR1 procedure but not the PR procedure (Nguyen et al., 2019). Equivalent breakpoints were also observed for the 0.06 and 0.15 mg/kg/infusion doses for the 1h-access, but not the 12h-access rats in a prior study (Wade et al., 2015), which suggests the current 6h-access produces a state more similar to the 1h than the 12h groups in that prior work.

The median split analysis for Cohort 1 confirmed that while a substantial mean difference in the number of infusions obtained was still present in the third week of acquisition, the percentage of responses made on the drug-associated manipulandum was identical across upper and lower halves of the distribution (**Figure S5**). A high, and equivalent level of this behavioral tuning was present throughout the course of the experiments, while the difference in oxycodone intake preference persisted through the fifth week. The ICSS pattern of increasing thresholds over the week also did not depend on average oxycodone intake across these preference sub-groups. Thus, these results are consistent with the fundamental principle of voluntary dug self-administration, i.e. that individual rats are responding for a subjectively equivalent level of intoxication.

The injection of THC (5 mg/kg, i.p.) elevated reward threshold (**Figure S7**), consistent with what is observed in naïve animals (Katsidoni et al., 2013; Vlachou et al., 2007); very low doses of THC (~0.1 mg/kg) are required to produce reward facilitation (decreased ICSS thresholds) in rats (Gardner et al., 1988; Katsidoni et al., 2013). Injection of THC elevated thresholds to a lesser extent when followed by a IVSA session, however this was not altered by the self-administration of vehicle versus oxycodone. Thus, it is most parsimonious to conclude that the additional hour since THC injection inherent to the post-IVSA ICSS assessment was the critical factor in the magnitude of the ICSS threshold change. There was a small reduction in threshold when oxycodone was self-administered for one hour under control conditions (e.g., when vehicle was administered prior to IVSA), and unchanged when saline was self-administered, as in our prior study (Nguyen et al., 2021) but this did not reach statistical significance.

**Differences with the Maguire and France Studies:** Related work from Maguire and France (Carey et al., 2023; Maguire and France, 2018, 2020) has been interpreted by the authors to show that cannabinoid agonists do not enhance the reinforcing value of a unit dose of an opioid. However, in their approach monkeys are assessed in four 25 minute sessions per day during which only 7 infusions are permitted. The remifentanil used in key studies is a very short acting opioid that does not appear to have a broad presence in human non-medical use; it is selected in animal work primarily because it leads to cocaine-like, stable self-administration patterns featuring high response rates. The authors' stated methodological intent to "limit drug accumulation" under low response requirements suggests that this approach does not let animals reach a self-selected satiety point to the extent expressed in our design. In addition, the JWH-018 and WIN compounds selected for some of the studies are full agonists at the $CB_1$ receptor and, as with remifentanil, such compounds have been typically less preferred over THC by most human users ever since the advent of synthetic cannabis products originally termed K2 or

Spice circa 2011 (Brents and Prather, 2014; Rosenbaum et al., 2012; Seely et al., 2012; Seely et al., 2013). These compounds also tend to have a shorter duration of action compared with THC. Further investigation would be required to integrate the two approaches and to determine if the different conclusions are a feature of the specific cannabinoid and opioid drugs, the design of the self-administration studies, the species used, or some combination of these factors.
